# MOPD I patient-derived cerebral organoids model microcephaly showing premature neurogenesis due to disrupted mitotic spindle orientation

**DOI:** 10.1101/2022.12.29.520610

**Authors:** Jagjit Singh, Noah J. Daniels, Filomena Pirozzi, Anthony Wynshaw-Boris, Rodrigo Lopez-Gonzalez, Richard A. Padgett

## Abstract

Mutations in the single-copy RNU4ATAC gene, which encodes U4atac snRNA of the minor spliceosome are linked to the developmental disorder microcephalic osteodysplastic primordial dwarfism type I (MOPD I). Partial loss-of-function mutations of U4atac snRNA lead to a poor prognosis, with less than three year survival. The most prominent characteristic of MOPD I is disrupted central nervous system development resulting in severe microcephaly and lissencephaly.

In this study, we used self-organizing 3D cerebral organoids from patient-derived induced pluripotent stem cells (iPSCs) to investigate defective cellular events that disturb the laminar organization of the cortex and influence brain topology. We analyzed organoids from iPSCs homozygous for the partial loss-of-function U4atac snRNA 51G>A mutation and compared them to isogenic organoids obtained from iPSCs expressing wild-type U4atac snRNA, using immunostaining and 10X Genomics single-cell RNA sequencing. In our MOPD I organoids, we observed: a) reduced proliferation accompanied by premature neurogenesis depleting the neuro-progenitor pool due to an increased frequency of horizontal cell divisions in the ventricular zone; b) reduced numbers of intermediate progenitor and outer radial glial cells in the outer sub-ventricular zone; and c) defective radial neuronal migration, which is critical for cortical expansion in humans. Our findings therefore provide insight into MOPD I cellular pathogenesis and underline the value of these cerebral organoids as model systems for human neurodevelopmental disorders.

## Introduction

Post-transcriptional removal of introns from the precursor mRNA is an important and tightly-regulated step in eukaryotic gene expression. The U12-dependent class of introns is a parallel subgroup that is distinct from the major U2-dependent introns. The U12-dependent class comprises <0.5% of the total introns in the human genome. A high degree of conservation in distantly-related organisms, occurrence in specific types of genes, and the slow rate of removal all indicate the significance of these low-abundance introns in gene regulation (1-3). The U12-dependent spliceosome requires a unique set of small nuclear RNAs (snRNAs) including U11, U12, U4atac, and U6atac, which are functional analogs of U1, U2, U4, and U6 in the U2-dependent spliceosome (4-6).

Several rare developmental disorders have been linked to biallelic mutations in the RNU4ATAC single copy gene that encodes the U4atac snRNA required for splicing the U12-dependent class of introns. These disorders include Roifman Syndrome (RS), Lowry-Wood Syndrome (LWS), and microcephalic osteodysplastic primordial dwarfism type I (MOPD I). These disorders have both distinct and overlapping symptoms and are inherited in an autosomal recessive pattern, where patients have biallelic homozygous or heterozygous point mutations in the single-copy RNU4ATAC gene (5-11). MOPD I can be particularly severe, with few patients surviving childhood, and it is associated with significant growth retardation (mean birthweight −5.8 SD, range −3.3 to −7.9 SD adjusted for gestational age), microcephaly (mean occipital frontal circumference (OFC) −7.0 SD, range −4.0 to −9.5 SD), and lissencephaly (7, 8). Other prominent MOPD I phenotypes include dry, aged-appearing skin, ridged cranial sutures, facial dysmorphism, and an average life expectancy of only 8.5 months (range 2.5 – 18 months) (12, 13). Defective U12-dependent splicing is lethal in Drosophila melanogaster, and results in developmental and growth defects in zebrafish embryos and Arabidopsis *thaliana*. Interestingly, U12-dependent intron splicing has been shown crucial for the survival of terminally differentiating retinal neurons in mouse (14-20).

The human brain cortex develops from a pseudostratified layer of neuroepithelial stem cells into a six-layered, functionally complex structure with multiple folds (gyrencephalic) on the surface. Cortical volume and folding are influenced by key cellular events, including expansion of the neuro-progenitor cell population (NPCs), differentiation of these NPCs into mature neurons, and neuronal migration (21-23). Genetic conditions or infectious diseases that interfere with these cellular processes trigger a number of cortical malformations, leading to mental retardation, mortality, and morbidity (24, 25).

A tightly-regulated balance between amplification of the neural progenitor cells and neurogenesis is required for proper development of the cortex. Overproduction of neurons at the expense of self-amplifying NPC divisions can deplete the NPC pool early in cortical development, resulting in microcephaly (26). The process of an NPC differentiating into a neuron is regulated by cell cycle length, particularly the lengths of G1 and mitosis (27-30). Cell-cycle defects that delay these phases result in increased neuronal differentiation. In addition, two types of NPCs, radial glial cells (RGCs) and intermediate progenitor cells (IPCs), contribute to neurogenesis. RGCs are required to maintain the NPC population, while IPCs primarily contribute to neuron production (31). In short, microcephaly can be caused by (a) death of the NPC or neuronal populations; (b) increased neuron production, resulting in depletion of the NPC pool; and/or (c) decreased IPC production, resulting in fewer neurons. Baumgartner et al. (2018) showed that the self-renewing radial glial cell population is sensitized towards the defects in U12-dependent splicing, leading to cell-cycle defects and apoptosis of cells causing microcephaly in U11 snRNA-ablated mice (20).

Miller-Dieker syndrome (MDS) is phenotypically similar but genetically different from MOPD I, as it is also characterized by the loss of cortical folding (lissencephaly) and is often associated with reduced brain size (microcephaly). Bershteyn et al. (2017), using MDS cerebral organoids as their model system, uncovered cell type-specific defects in lissencephaly spanning the stages of neuroepithelial cell expansion and neuronal migration, as well as the mitotic properties of outer-radial glial (oRG) progenitors, providing deeper insight into human cortical development and lissencephaly (32).

Severe MOPD I due to homozygous *RNU4ATAC* 51G>A mutations is found in the Ohio Amish population. We have previously investigated the effects on U12-dependent intron splicing using cell lines derived from these patients. We found that the mutant U4atac snRNA was defective for protein binding, leading to lower levels of the essential U4atac/U6atac U5 tri-snRNP complex. We further showed that expression of wild-type U4atac snRNA, using lentiviral delivery, restored minor intron splicing to normal levels in induced pluripotential stem cells (iPSCs) obtained by reprograming of umbilical cord fibroblasts (33).

The mouse brain is lissencephalic in nature; therefore, there are certain aspects of human cortical development that may not be adequately recapitulated in mice; thus three-dimensional cerebral organoids derived from human induced pluripotent stem cells have been used to this aim, as they possess a more similar cellular heterogeneity to the human cortex (34-38). Here, we use MOPD I patient iPSC-derived cerebral organoids to investigate possible defects in cerebral development that may contribute to the microcephaly and lissencephaly that characterize this disease. Specifically, we show that patient-derived organoids have overall growth defects and show enhanced neurogenesis that leads to a depletion of the pool of NPCs. We also find that neurofiber outgrowth and cell migration are defective in the mutant organoids compared to isogenic control cells.

## Results

To understand the underlying mechanisms that lead to the neurological structural defects seen in MOPD I patients, we used brain organoids from previously published patient-derived iPSCs with homozygous 51G>A mutations in U4atac snRNA (33). For comparison, we generated isogenic cells lines using lentiviral delivery to introduce either an empty vector construct (cell line herein denoted as MOPD I) or a construct expressing a wild-type RNU4ATAC gene as previously described (33) (herein denoted as MOPD I Rescued) (Supp. Figure 1A). These isogenic iPSCs grew as colonies and showed similar doubling rates (Supp. Figure 1B). Bulk RNA-seq of both the MOPD I iPSCs and the MOPD I-Rescued cell lines showed robust expression of pluripotential marker genes (MYC, SOX2, OCT4, and NANOG) and low expression of neural genes prior to differentiation (EOMES, SOX1, PAX6, SATB2, CTIP2, and MAP2) (Supp. Figure 1C). These iPSCs mimic the splicing defects of U12-dependent introns similar to the original patient fibroblasts, and this defect could be rescued by the ectopic expression of WT U4atac (33). Consistent with Jafarifar et al (2014), our bulk-RNA sequencing analysis show increased frequency of U12-dependent intron retention in MOPD I iPSCs when compared with MOPD I-Rescued cells (Supp. Table 1) (2, 33).

### MOPD I organoids recapitulate microcephaly and cortical layering defects *in vitro*

To generate brain organoids, we followed a modified version of the Sasai protocol (39) (Figure 1A). All iPSC lines tested generated spheroids and differentiated into forebrain organoids. The size of the organoids was monitored during the organoid differentiation process; at day 24, a difference in organoid size between MOPD I and MOPD I-Rescued became evident (Figure 1B and Supp. Figure 2). We quantified organoid area at two time points, 24 days and 32 days of differentiation, and found that MOPD I organoids were reduced (∼2.5 fold median value) in size compared to rescued organoids (Figure 1C). Interestingly, the surface of our MOPD I organoids presented fewer lobes and protrusions than the MOPD I rescued organoids, suggesting defects in radial neuronal migration (Supp. Figure 2A and 2B).

**Figure 1.**
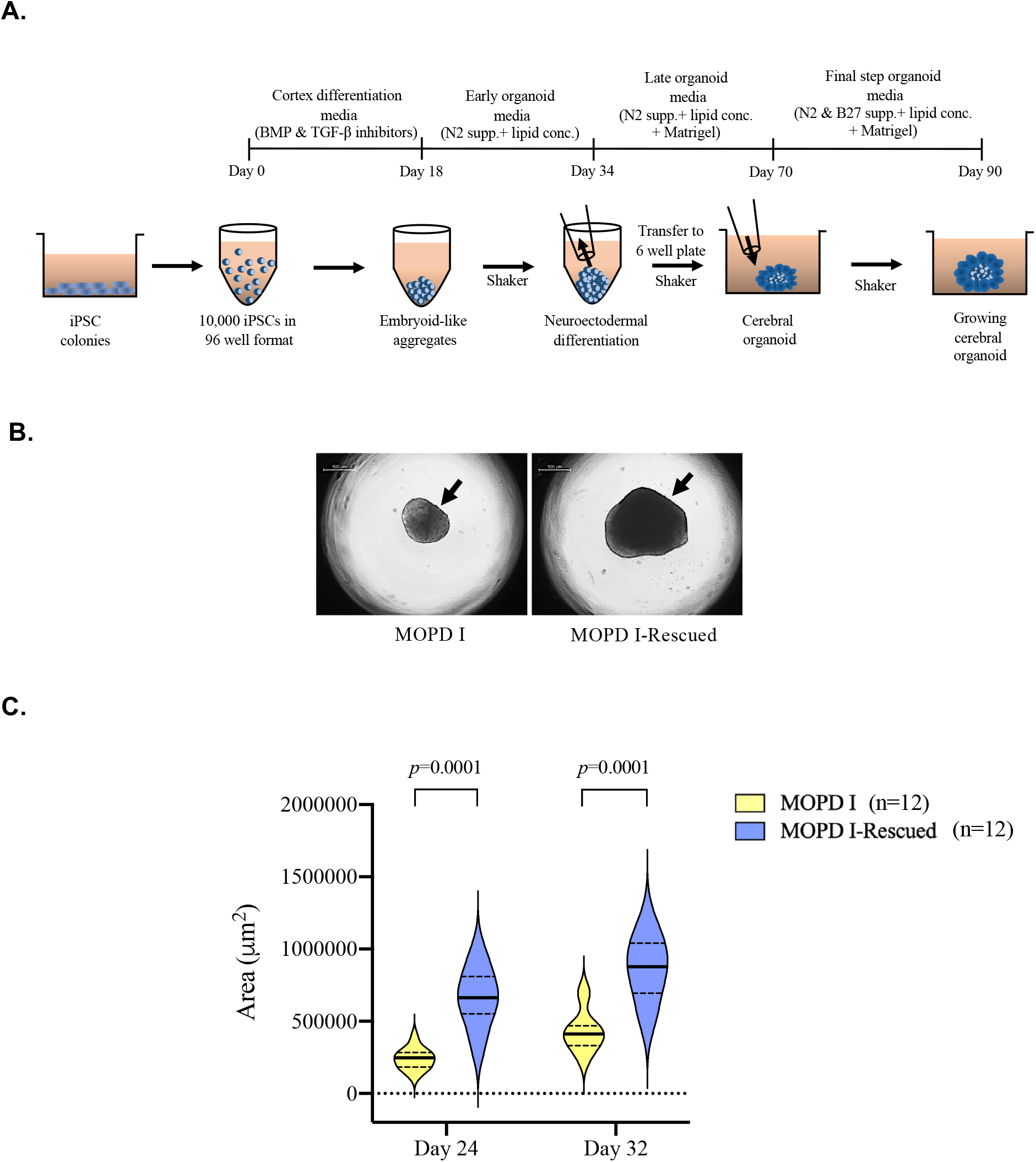
MOPD I Organoids Recapitulate Microcephaly. A. Schematic diagram of forebrain organoid differentiation from iPS cells. The timeline shows the differentiation steps and media composition from day 0-90. B. 3-D cerebral organoids were generated from MOPD I and MOPD I-Rescued iPS cells. Representative images of organoids at 24 days showed MOPD I organoids were smaller in size compared to the MOPD I–Rescued organoids. Scale bar, 500µm. C. Forebrain organoids (MOPD I and MOPD I-Rescued) were monitored for size at different time points. Quantification of area was performed using ImageJ. Violin plots show smaller sized MOPD I organoids compared to MOPD I-Rescued organoids at both day 24 and 32. *p*-values were computed using unpaired student’s-t test.

Once we established a significant difference in size between rescued and MOPD I organoids, we decided to analyze their cell type composition. We fixed, cryo-sectioned and immuno-stained organoids at 60 days of differentiation to assess the presence and distribution of specific neuronal cytotypes, as well as cell cycle markers. At this time point, the organoids consisted of NPCs organized into multiple pseudostratified ventricular zone (VZ)-like regions (Figure 2A). Immuno-staining and further quantification revealed reduced (∼4.0 fold median value) numbers of TBR2-positive intermediate progenitor cells and increased (∼2.0 fold median value) numbers of MAP2-positive neurons in MOPD I organoids as compared to MOPD I-Rescued organoids (Figure 2B and 2C).

**Figure 2.**
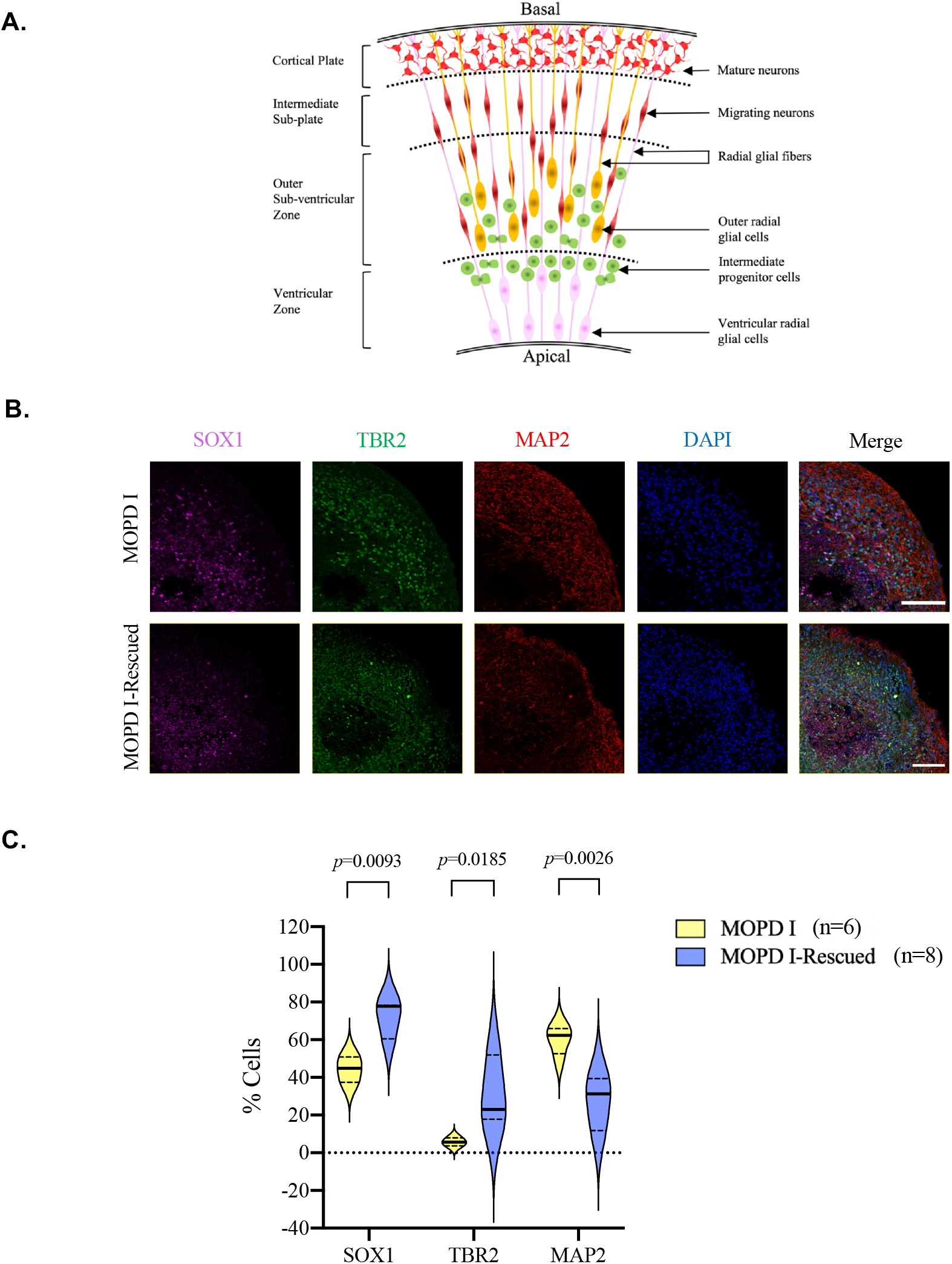
MOPD I organoids showed increased neurogenesis depleting the neuro-progenitor pool. A) Schematic representation of different cell types that form the laminar organization in the human brain cortex. Ventricular radial glial cells (pink), intermediate progenitor cells (green), outer radial glial cells (yellow), migrating neurons in the subcortical plate and mature neurons in the cortical plate (both in red) are organized in an apico-basal directional manner. B) Immunostaining of forebrain organoids at 60 days of differentiation. The ventricular radial glial cells are probed for SOX1 labeled in pink, the intermediate progenitor cells are probed for TBR2 labeled in green and mature cortical neurons are probed for MAP2 labeled in red. DAPI was used to stain the nuclei. Scale bar 100 µm. C) Quantification of SOX1+, TBR2+ and MAP2+ cells in MOPD I and MOPD I-Rescued organoids after 60 days of differentiation. We use QuPath to compare the percentage of positive cells in the two different conditions. Two-way ANOVA followed by Bonferroni’s adjustment for multiple comparisons was performed to determine statistical significance.

Consistent with their reduced size, MOPD I organoids showed reduced (∼5.0 fold median value) proliferation in the VZ-like regions, based upon the number of Ki67-positive cells in these areas (Figure 3A and 3B). When we analyzed the number and distribution of SOX1+ cells in MOPD I organoids, we found a reduced number of cells and a homogeneous distribution throughout the organoid, in contrast to a more organized and localized presence of SOX1+ cells in the VZ-like areas of rescued organoids. Interestingly, there was no significant difference in the number of apoptotic cells in the MOPD I organoids when compared to the MOPD I-Rescued (Figure 3C and D). These results suggest that the MOPD I organoids undergo premature neuronal differentiation leading to depletion of the neural progenitor cell pool, rather than increased apoptosis of NPCs or neurons.

**Figure 3.**
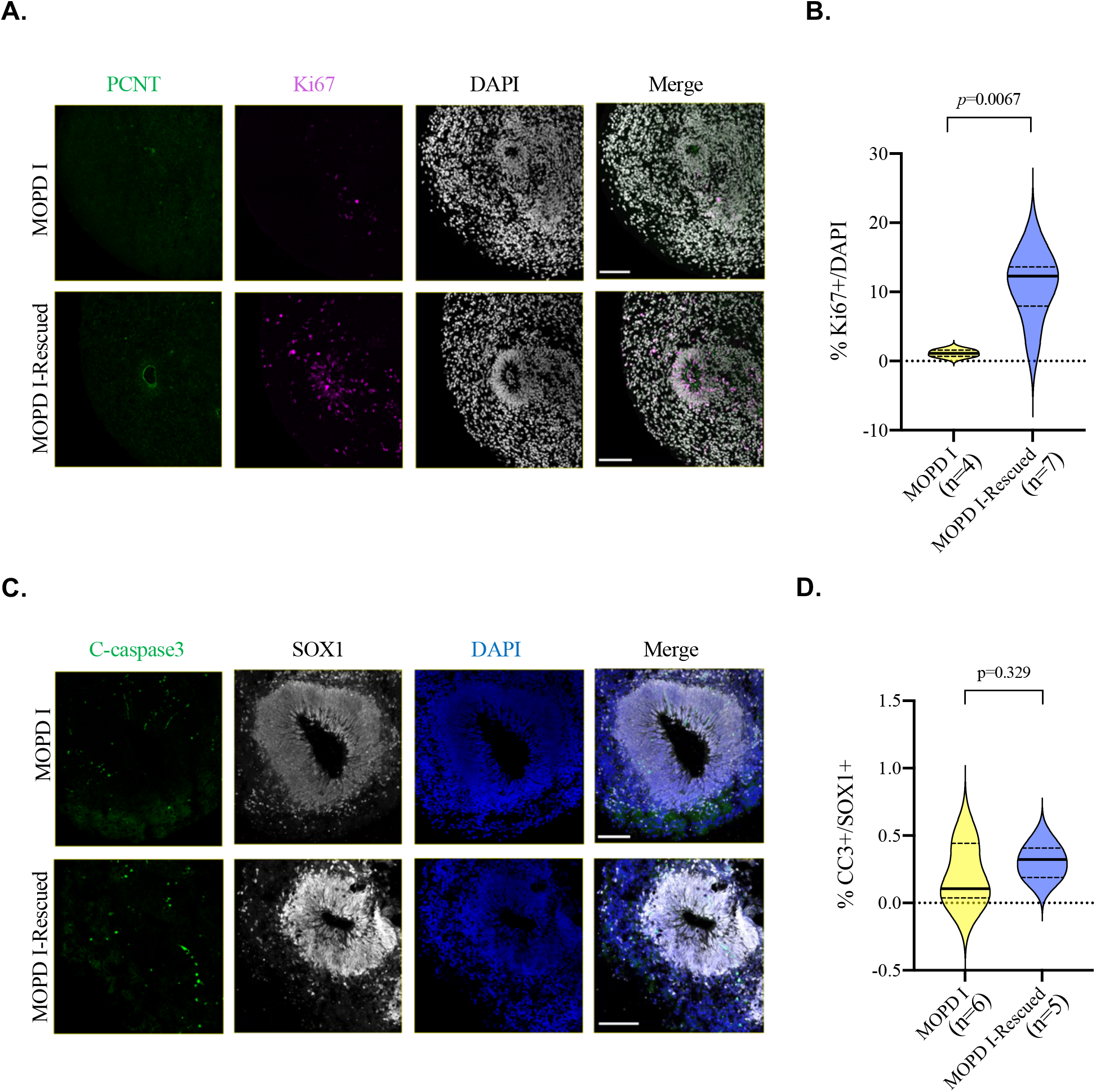
Reduced Proliferation of neural progenitor cells in MOPD I organoids. A. Representative confocal images of immunofluorescent staining of the MOPD I and MOPD I-Rescued organoids probed for Ki67 and PCNT. Scale bar 100 µm. B. Percentage of Ki67+ cells in MOPD I and MOPD I-Rescued organoids at 60 days of differentiation. We used QuPath for quantification. Statistical analysis was done using Mann-Whitney unpaired t test. C. Representative confocal images of immunofluorescent staining of MOPD I and MOPD I-Rescued organoids probed for neuronal progenitors in the ventricular zone (SOX1) and for active apoptosis (cleaved Caspase 3). Scale bar 100 µm. D. Quantification of SOX1+ cells in VZ-like regions expressing cleaved-caspase 3 was performed using QuPath. A Mann-Whitney unpaired t-test was performed to assess statistical significance.

### Increased frequency of horizontal plane of division in the VZ-like regions of MOPD I organoids

Regulated orientation of the mitotic spindle has been shown to play a very important role in determining cell fate decisions during development in a variety of systems. Previous studies show that during the early stages of development of the neocortex, the vertical plane of division, which produces symmetric NPCs, is very important as it populates the neural progenitor pool in the extended VZ (Figure 4A). The switch from the vertical plane of division to the horizontal leads to premature neurogenesis, which depletes the neural progenitor pool and has been associated with cortical thinning and microcephaly in other systems (40-46). To determine whether a similar defect occurred in MOPD I organoids, we immunostained MOPD I and MOPD I-Rescued organoids at day 40 of differentiation using the specific mitotic marker p-Histone-H3 and Pericentrin (PCNT) to identify the apical surface. We found a significant increase in the frequency of cells presenting a horizontal plane of division when compared to the MOPD I-Rescued organoids (Figure 4B and 4C). This suggests that the increase of MAP2 neurons in MOPD I organoids may be the result of a switch in division plane in NPCs.

**Figure 4.**
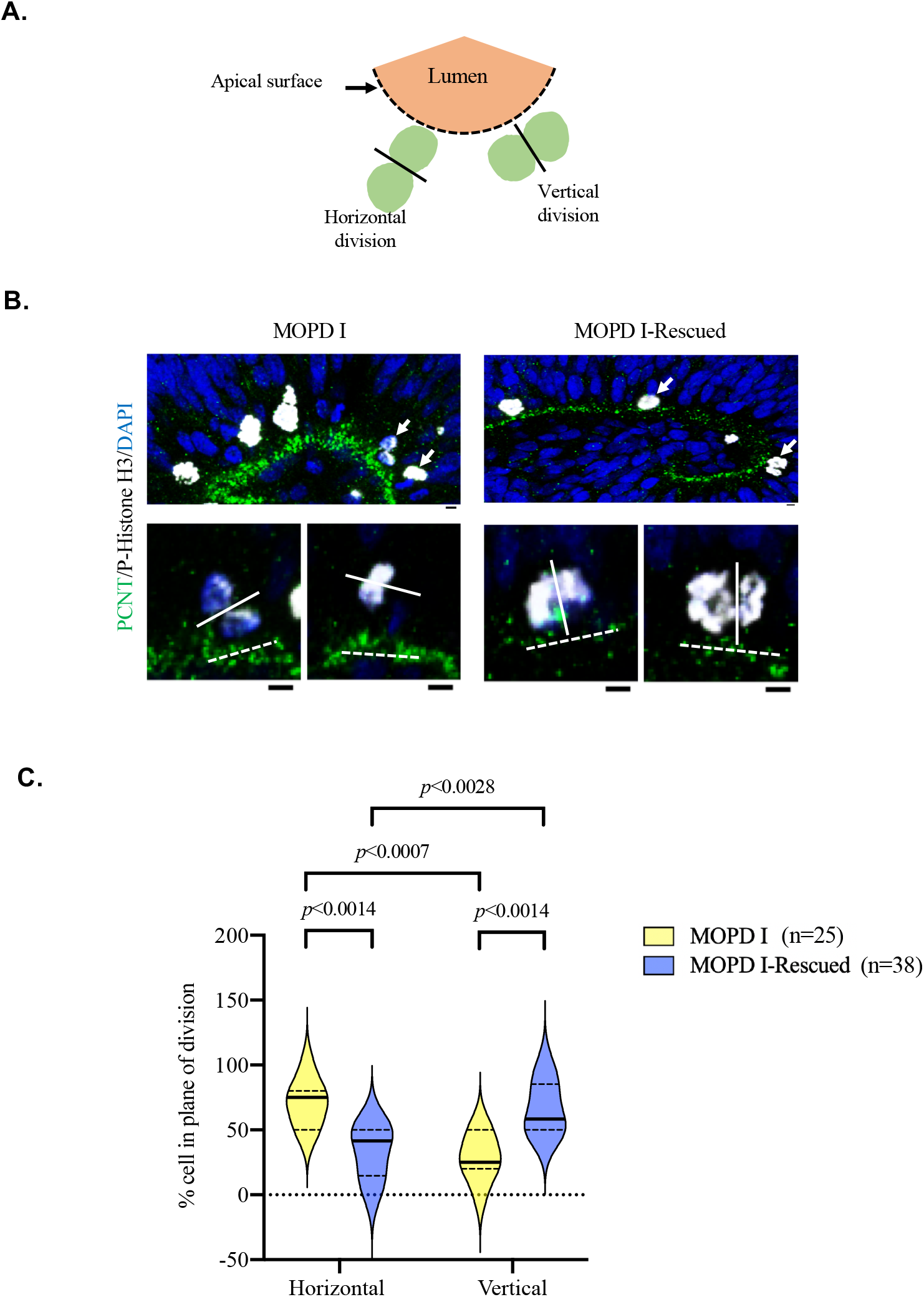
Increased frequency of horizontal nuclear division in MOPD I organoids. A. Schematic representation of neural progenitor cells in the ventricular zone undergoing two different types of nuclear divisions (vertical and horizontal) with respect to the apical surface. The dotted line represents the apical surface around the fluid filled lumen/ventricle. B. Sections of 60 days cultured MOPD I (left panel) and MOPD I-Rescued (right panel) organoids were stained with PCNT, Phospho-histone H3 antibodies and DAPI to assess the plane of division at the apical surface. Scale bar 100 µm. C. Quantification of the percentage of nuclei undergoing a particular plane of division, where (n) is equal to the total number of dividing nuclei measured. Two-way ANOVA followed by Bonferroni’s adjustment for multiple comparisons was performed to determine statistical significance.

### MOPD I organoids show migration defects of deep layer neurons and reduced numbers of outer radial glial cells

In order to analyze cortical patterning, we analyzed co-expression of the dorsal telencephalon progenitor marker SOX1 in the VZ-like regions, which are surrounded by deep-layer subcortical projection neurons positive for CTIP2 and SATB2 (47, 48). Immunostaining and further quantification showed no differences in the numbers of CTIP2+ and SATB2+ neurons between MOPD I and MOPD I-Rescued organoids (Figure 5A and 5B). However, when we analyzed the number of outer radial glial cells, we found a significant reduction (∼2.0 fold median value) in HOPX+ cells in MOPD I organoids as compared to MOPD I-Rescued organoids (Figure 5C and 5D), suggesting that one of the reasons behind radial neuronal migration defects could be a reduced number of radial neuronal fibers that are used as the substrate for migration.

**Figure 5.**
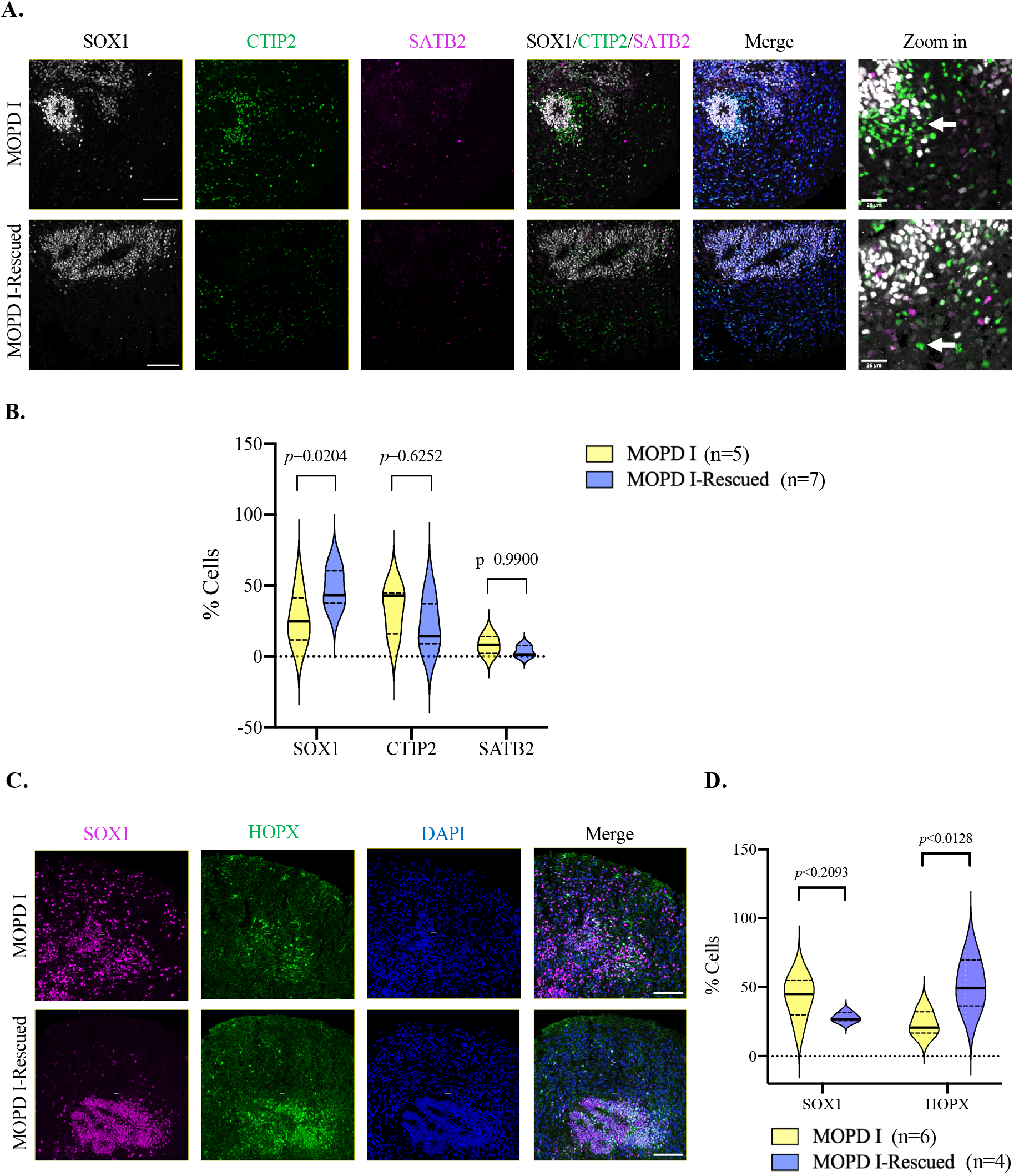
MOPD I organoids show migration defects of deep layer neurons and a reduced number of outer radial glial cells. A. Immunostaining of forebrain organoids at 60 days of differentiation. The ventricular radial glial cells are probed for SOX1 labeled in white, the deep layer neurons (early neurons) are probed for CTIP2 labeled in green and the upper layer neurons are probed for SATB2 (late neurons) labeled in magenta. DAPI was used to counterstain the nuclei. Scale bar 100 µm. Zoomed-in image with scale bar 25 µm. B. Quantification of the percentage of SOX1+, CTIP2+ and SATB2+ cells in MOPD I and MOPD I-Rescued organoids. Two-way ANOVA followed by Bonferroni’s adjustment for multiple comparisons was performed to determine statistical significance. C. Immunostaining of forebrain organoids at 60 days of differentiation. The ventricular neuronal progenitor cells are probed for SOX1 (in magenta) and the outer radial glial cells are probed for HOPX (in green). DAPI was used to counterstain the nuclei. Scale bar 100 µm. D. Quantification of the percentage of SOX1+ and HOPX cells in MOPD I and MOPD I-Rescued organoids. Two-way ANOVA followed by Bonferroni’s adjustment for multiple comparisons was performed to determine statistical significance.

### MOPD I organoids showed defective radial neuronal migration

Defective radial migration of cortical neurons is a well-established developmental phenotype in lissencephaly (32, 49-54).We performed a migration assay modified from a previously published protocol in Schaffer AE et al (2018) using our human organoid model system (55). Briefly, we grew organoids from MOPD I and MOPD I-Rescued cell lines for 35 days and then embedded the live organoids in concentrated matrigel in a glass-bottom plate. After 24 hours of adhesion, the organoids produced numerous neuronal processes extending from the edge of the organoids into the matrigel (Figure 6A). These processes showed prominent expression of DCX and TUJ1, suggesting that they represent bundles of neurites and radial glial fibers. We observed fewer of these processes in the MOPD I organoids as compared to the MOPD I-Rescued organoids. After 72 hours, DAPI-stained cells were observed migrating radially outward along these processes (Figure 6B). We measured the placement of neuronal nuclei from the edge of the organoid at 72 hours. The quantification of the neuronal displacement in MOPD I organoids show that the majority of the neurons failed to migrate beyond the organoid edge (Figure 6C and 6D). In our MOPD I organoids ∼80% of the neurons migrated in the range of 0-200µm, ∼5% were in the range of 201-400µm, ∼2.5% in 401-600µm and ∼2.5% migrated to the 601-800µm range. Whereas, MOPD I-Rescued organoids showed uniform migration of neurons throughout 0-800µm distance range. ∼30% of the neurons were in the distance range of 0-200µm, ∼40% of the neurons were on 201-400µm, ∼20% neurons migrated to 401-600µm and ∼10% migrated neurons were in the range of 601-800µm (Figure 6E).

**Figure 6.**
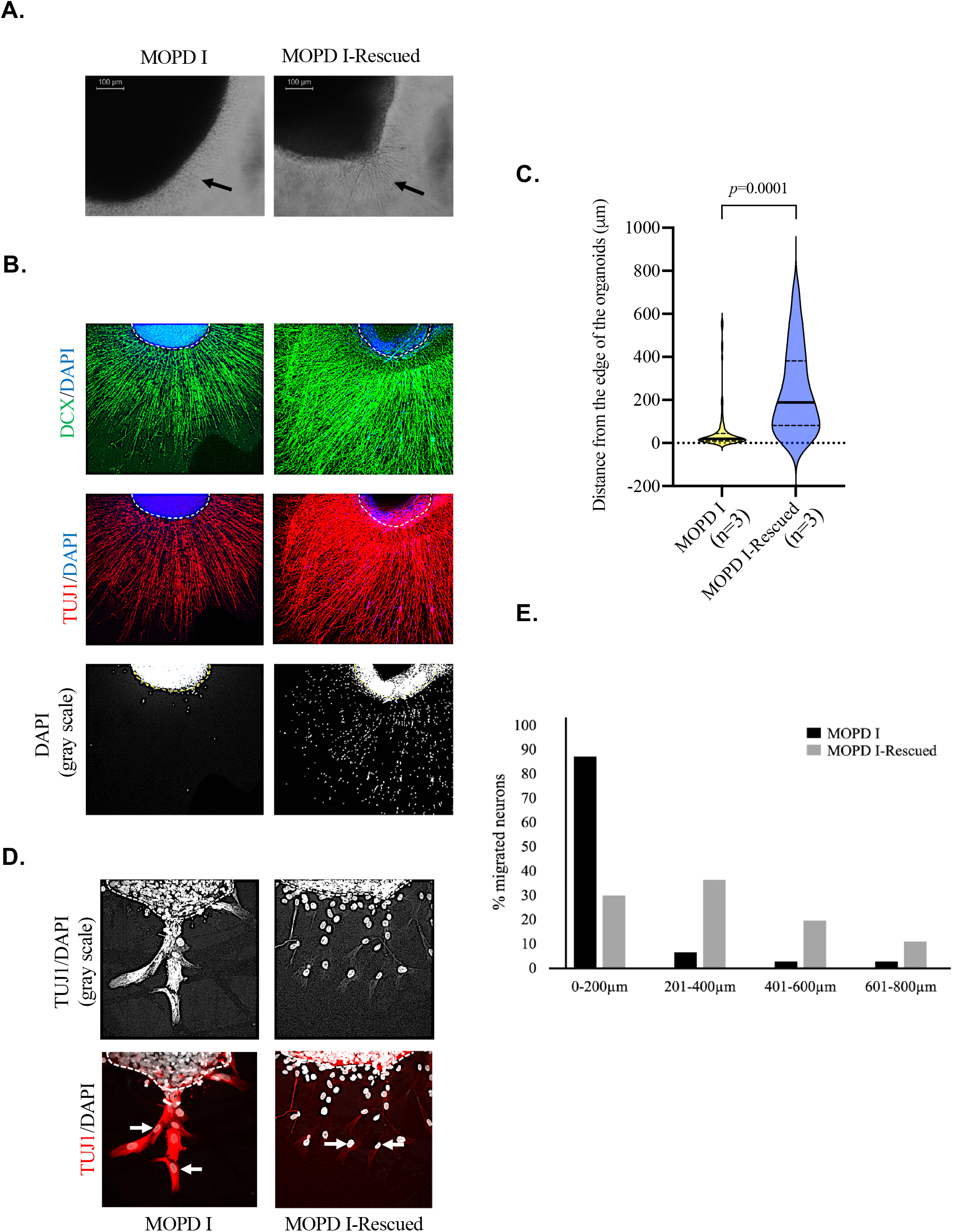
Defective neuronal migration along radial glial fibers in MOPD I organoids. A. Organoids from MOPD I and MOPD I-Rescued cell lines were cultured for 35 days and were embedded in concentrated matrigel. After 24 hours, neuronal processes (indicated by the black arrows) were observed growing into the matrigel. Scale bar 100 µm. B. Immunostaining of MOPD I and MOPD I-Rescued organoids 72 hours post embedding. Organoids were fixed and stained with DCX (green), Tubulin-Beta III/TUJ1 (Red) antibodies and DAPI. C. Quantification of the distance of migrating neurons in MOPD I organoids as compared to MOPD I-Rescued. Mann-Whitney unpaired t-test was performed to determine statistical significance. N represent the number of organoids tested for each group.

### MOPD I organoids show premature differentiation and decreased proliferation using scRNA Sequencing

In order to support and extend our immunohistochemistry analyses, we analyzed single-cell gene expression patterns from MOPD I and MOPD I-Rescued organoids after 10 weeks of differentiation. Briefly, we enzymatically dissociated our 70-day cultured live organoids into a single-cell suspension (see STAR methods). Cell viability was above 90%, with 10,000 cells per cell line captured by the 10X Genomics system. Following quality control, we clustered 4,448 and 5,006 cells for MOPD I and MOPD I-Rescued organoids, respectively (Supp. Figure 3A). We identified 6 major cell types: neurons, radial glia, choroid plexus, cycling progenitors, intermediate progenitors, and unfolded-protein-response-related cells (UPRC) (Figure 7A-C, Supp. Figure 3B). Comparison of cell type composition revealed a decrease in the proportion of radial glia, cycling progenitors, intermediate progenitors, and choroid plexus in addition to an increase in neurons in MOPD I organoids compared to MOPD I-Rescued (Figure 7D).

**Figure 7.**
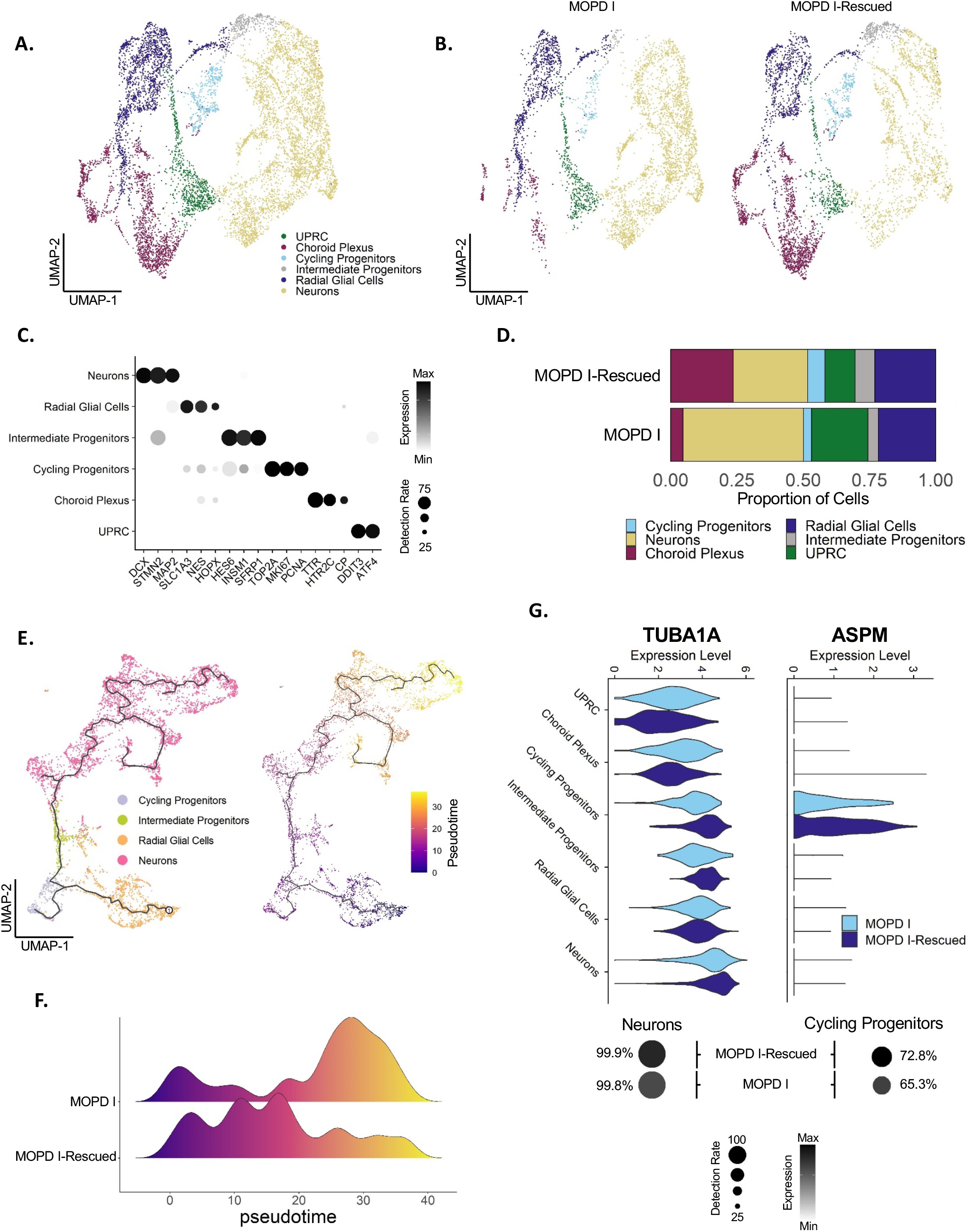
Single cell RNA-seq of organoids shows clustering of different cell populations revealing increased neurogenesis and reduced number of neuronal progenitor cells. A) UMAP of combined organoids color-coded by the six main clusters associated with specific cell types: UPRC, unfolded-protein-response-related-cells (green); Choroid Plexus (red); Cycling progenitors (light blue); Intermediate Progenitors (grey); Radial Glial Cells (dark blue); and Neurons (yellow). B) UMAP of MOPD I and MOPD I-Rescued organoids. Cells are color-coded by cell type following the same legend as in panel A. MOPD I=4448 cells; MOPD I-Rescued=5006 cells. C) Expression levels of genes used to identify cell types. Dot size shows percentage of positive cells. D) Bar plots indicate proportion of cell types in each organoid group. E) Developmental trajectory of neuronal differentiation using Monocle3. UMAP of cell types (left) and pseudotimes (right) using radial glial cells as the starting node. F) Distribution of pseudotimes along neuronal differentiation for MOPD I and MOPD I-Rescued. Two-sided Kolmogorov-Smirnov test. P=2.2e-16. G) Cell type specific expression of TUBA1A and ASPM in MOPD I and MOPD I-Rescued organoids (top panels). Relative expression level and percentage of cells expressing TUBA1A in neurons (bottom left) and ASPM in cycling progenitors (bottom right). Dot size shows percentage of positive cells.

The decrease in intermediate progenitors and increase in neurons in MOPD I organoids suggests that the expansion of neurons could be due to premature neurogenesis and accelerated neuronal differentiation. To investigate this, we generated developmental trajectories of our cerebral organoids using Monocle3 (Supp. Figure 3C). Our organoids progressed from radial glia cells bifurcating to choroid plexus or neuronal cell types (Figure 7E). To gain a better understanding of the neuronal differentiation timeline, we subclustered cells progressing from radial glia to mature neurons and performed pseudotime trajectory analysis (Figure 7E, Supp. Figure 3D). Interestingly, the developmental trajectory was significantly skewed towards the mature neuronal population for the MOPD I organoids, while MOPD I-Rescued cells predominantly clustered at intermediate progenitor and early neuronal stages (Figure 7F, pvalue < 2.2–16). These data are in agreement with the interpretation of premature neurogenesis that we found in the immunostaining analysis (Figure 2B and 2C; Supp. Figure 3E).

We next explored the cell type-specific and molecular changes in our organoids by performing differential expression analysis for each cell type-specific cluster between our MOPD I and MOPD I-Rescued organoids. Intermediate progenitors had the highest number of differentially-expressed genes (DEGs; Supp. Figure 4A). Overall, we found more downregulation of DEGs than upregulation for all cell type-specific clusters besides neurons. Additionally, most of the gene dysregulation was specific to each cell type-specific cluster, with minimal overlap of DEGs (Supp. Figure 4B). To gain a better understanding of dysregulated pathways, we performed Gene Ontology (GO) analysis for our clusters progressing through neuronal development (Supp. Figure 4C). Strikingly, we observed an upregulation of genes involved in neuronal migration in our cycling progenitor population in the MOPD I-Rescued organoids (Supp. Figure 4C and 5). Additionally, cerebellar development genes, such as NEUROD1 and NEUROD2, were upregulated in neurons from our MOPD I-Rescued organoids (Supp. Figure 4C and 5). These data were further supported by weighted gene co-expression network analysis (WGCNA). We observed an upregulation of genes involved in neuronal development for intermediate progenitors and early neurons, in addition to a decrease in neuronal maturation gene expression for our MOPD I-Rescued organoids. These data support a premature neurogenesis phenotype with defects in neuronal migration (Supp. Figure 3E).

Of note, differential gene expression analysis from the 10X Genomics single-cell RNA sequencing data showed that MOPD I organoids presented reduced expression of genes ASPM and microtubule organization TUBA1A (Figure 7G). Reduced expression of these genes has been previously associated with lissencephaly and neurodevelopmental disorders, and involved in regulation of mitotic spindle orientation (56-60).

## Discussion

In this work, we used previously-characterized iPSC lines to generate brain organoids and investigate pathological mechanisms of MOPD I mutations (33). Analyses of forebrain organoids from MOPD I patients showed reduced proliferation, premature neuronal differentiation, increased horizontal cell divisions in VZ-like areas, and reduced radial glial migration. Patient-derived iPSCs offer an opportunity to uncover molecular mechanisms behind microcephaly, as they have been successfully used to model human neurological disorders *in vitro* since they possess the same genetic make-up of the individual that they were derived from (61-73). In addition, modeling these neurodevelopmental disorders in mice is not ideal since the mouse brain is lissencephalic in nature. Moreover, comparison of the laminar organization of rodent cortex and humans revealed a lack of outer radial glial cells, which form an additional layer of cells in the cortex called the outer sub-ventricular zone (74). This layer of cells is responsible for the complexity and 3-fold expansion of the cortex in humans. For this reason, mouse models fail to fully recapitulate microcephaly in humans.

Our analysis of MOPD I organoids showed decreased proliferation of SOX1+ NPCs in the VZ-like areas, confirming the hypothesis that depletion of the NPC pool might be a leading cause of the reduced brain size in these patients. In addition, MOPD I organoids showed increased numbers of cells expressing early neuronal markers, both at the immunohistochemistry level as a significant increase in MAP2+ neurons, and at the single-cell RNA analysis level showing an increase in neurons and a reduction in radial glia, choroid plexus, and cycling and intermediate progenitors, adding premature neuronal differentiation as a parallel mechanism for the observed microcephaly. A similar phenotype was previously described in a brain organoid model of microcephaly caused by genes other than RNU4ATAC (32, 73, 75).

Furthermore, to determine whether the reduction in size of MOPD I organoids was due to cell death, we analyzed the proportion SOX1+ cells that were positive for cleaved caspase 3. We found no differences between MOPD I and MOPD I-Rescued organoids, suggesting that apoptotic cell death does not contribute to the reduction in size in MOPD I brain organoids.

During development of the brain cortex, before a cell prepares to divide, the intracellular components must be partitioned into two daughter cells. The cells can undergo symmetric or asymmetric mitotic division. Symmetric division ensures identical distribution of cellular components between the two daughter cells. In asymmetric division, components such as cell fate regulators may be partitioned asymmetrically to result in different cell fates following division (76). Asymmetric division results in the generation of many distinct cell populations. The asymmetric distribution of these subcellular components can be achieved by regulating mitotic spindle orientation (77, 78). Dysregulation of mitotic spindle orientation has previously been associated with microcephaly *in vivo* and *in vitro*, as in the cases of mutations in ASPM, WDR62, CDK5RAP2, and LIS1 (56-58, 66, 79-83). In line with these findings, our results reveal that MOPD I organoids present an increased number of NPCs following an asymmetric (horizontal) division when compared to MOPD I-Rescued organoids. The evidence of increased frequency of horizontal plane of nuclear division in MOPD I organoids supports an interpretation of premature neurogenesis. This mechanism is shared with another recent cerebral organoid model of lissencephaly and microcephaly due to defects in mitotic spindle orientation (32), highlighting how changes in the angle of the division plane can deeply impact neuronal differentiation (Figure 8).

**Figure 8-Model.**
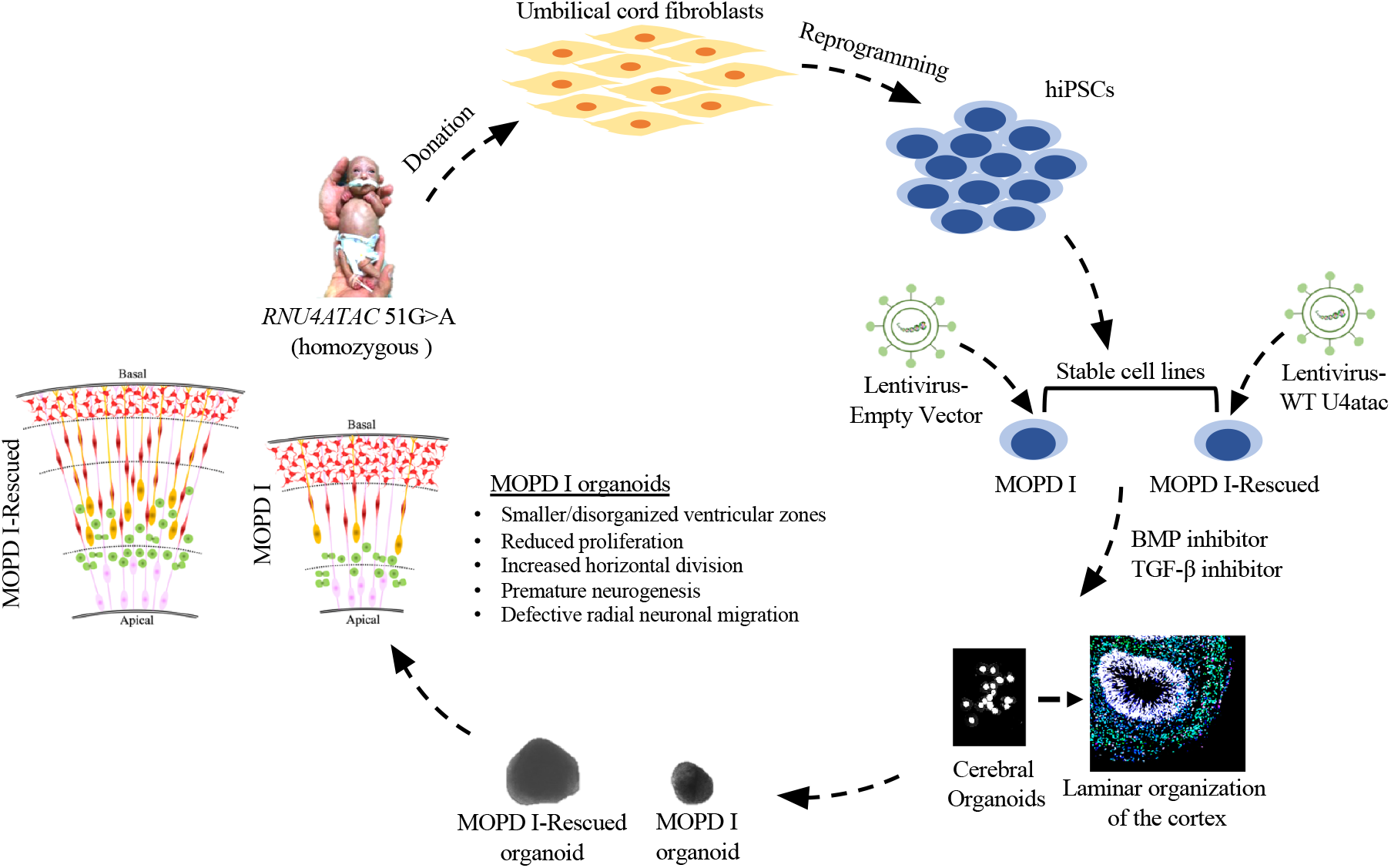
Illustration of MOPD I organoids recapitulating microcephaly. Cortical organoids from MOPD I patient-derived iPSCs model microcephaly. Reduced proliferation and increased frequency of horizontal divisions are observed in neuro-progenitors cells in MOPD I organoids. Cortical neurons are more abundant in MOPD I organoids. Reduced outer radial glial cells contribute in neuronal migration defects.

Radial glia fiber are crucial for proper migration of newly formed neurons to their final cortical destination, as described previously (84-86). Defects in radial migration of cortical neurons have been established as developmental phenotypes that contribute to lissencephaly (50-55). Here, we found that MOPD I-Rescued organoids projected significantly more DCX+ and TUJ1+ neurites and radial glial fibers as compared to MOPD I organoids. In addition, we found a reduced number of migrating nuclei from MOPD I as compared to MOPD I-Rescued organoids in our migration assay. This suggests that MOPD I organoids also exhibit a correlation between the reduced expression of TUBA1A in cell populations like cycling progenitors, intermediate progenitors and neurons and the defective neuronal migration (Figure 8).

Our single-cell RNA-seq analysis further validates our immunohistochemical results. Our identification of different cell populations and further comparison between MOPD I and MOPD I-Rescued organoids revealed a decrease in the proportion of radial glia, cycling progenitors, intermediate progenitors, and choroid plexus cells in addition to an increase in neurons for MOPD I. Moreover, MOPD I organoids revealed decreased expression of genes previously associated with microcephaly such as ASPM (87), RELN (88), and TUBA1A (89, 90). All of these genes have been reported as crucial for both radial and tangential neuronal migration in the cortex, as well as for the modulation of neuronal migration, explaining the migration defects observed in MOPD I organoids. In conclusion, by using human iPSC-derived forebrain organoids as a model system, we provide new insights into the pathogenesis of MOPD I and a robust working model that will be used to test new potential therapeutic strategies.

## Materials and Methods

### iPSC reprograming and culture methods

The umbilical cord fibroblasts from two MOPD I patients with the 51G>A mutation were gifts from Dr. A. de la Chapelle’s laboratory (7). iPSC cell lines were derived from a 51G>A MOPD I patient and control umbilical cord fibroblasts at the pluripotent stem cell facility at Case Western Reserve University using retroviral transduction of OCT4, KLF4, SOX2, and C-MYC as previously described (91). Results from the characterization of these cell lines were previously published (33). The iPSC lines were cultured on feeder-free Matrigel-coated plates in mTeSR media (StemCell Technologies) before differentiation into forebrain cerebral organoids.

### Generation of isogenic iPSC cell lines

Lentiviruses carrying wild-type U4atac and Empty Vectors (separately) were obtained by transfecting HEK 293T cells (ATCC, ACS-4500) using Lipofectamine 2000 in FBS free Optimem together with the virus packaging plasmids (psPAX2 and pMD2-G, Addgene # 12260 and # 12259) and the plasmid expressing either wild-type U4atac or an Empty Vector. The plates were incubated at 37°C/ 5% CO_2_ for 3 hours. 7 ml of DMEM supplemented with FBS was added to the previous transfection media on the plate and the plates were incubated at 37°C overnight. The virus stocks were harvested at 24h and 48h after transfection, collecting and centrifuging the media at 300g for 4 mins. The stock was stored at -20°C. Ten ml of fresh media was added and the plates were incubated at 37°C/ 5% CO_2_ overnight for the next virus harvest. Similarly, the second virus stocks were collected the next day. iPSC lines at 50% confluency were infected using 1 ml of viral stocks in a 6 well-plate format for 24 hours at 37°C. The media was aspirated and the viral infection was performed again after 24 hours. After 24 hours, the cells were exposed to puromycin (0.5μg/ml)for selection of the population of interest. The selected cells were further cultured for the downstream experiments.

### Generation of cerebral organoids and culture

We selected the Sasai protocol (with some modifications) to differentiate our iPSC lines into organoids, as it allows for increased heterogeneity in the neuronal cell types obtained because it is a self-patterning protocol (38). Briefly, iPSCs were dissociated into single cells with 1mL Accutase and re-aggregated using cortical differentiation medium in lipidure-coated 96-well V-bottom plates (Sbio, #MS9096VZ) at a density of 1x10^4^ cells per well, 100μl per well. The cortical differentiation medium (Glasgow-MEM, 20% KSR, 0.1mM NEAA, 1mM sodium pyruvate, 0.1mM beta-mercaptoethanol, 100 U/mL penicillin and 100U/ml streptomycin) was supplemented with Rho Kinase inhibitor (Y-27632, 20 mM, days 0-3), SMAD inhibitor (LDN 200nM, days 0-3), and TGF-b inhibitor (SB431542, 10 μM, days 0-3). When the spheroids were formed, the cortex differentiation media was prepared without Rho Kinase inhibitor and the concentrations of SMAD inhibitor and TGF-b inhibitor were reduced by half (from day 4 onwards). Media changes were performed on day 3 and then every other day until day 18, and images were taken by using an inverted microscope (Leica DM IL # 090-135.002; Leica EC3) for monitoring the size of the organoids. On day 18, the aggregates were transferred (20 organoids/well) to ultra-low adhesion 6-well plates with Early organoid media DMEM/F12 medium with Glutamax supplemented with N2, Lipid Concentrate, Fungizone (2.5 mg/mL), and penicillin/streptomycin (100 U/mL) and grown under 40% O_2_/5% CO_2_ conditions. Late organoid media was used to culture the organoids from day 35 until day 55, supplemented with FBS (10% v/v), Matrigel (1% v/v) and heparin (5 mg/mL). Final step organoid medium supplemented with double the concentration (2%) of Matrigel and B27 supplement was used to culture the growing organoids from day 57 until day 70. Organoids were fixed with 4% paraformaldehyde in PBS for 30 min, washed with PBS, and dehydrated with 30% sucrose in PBS overnight at 4°C. After the complete dehydration, organoids settled in the bottom of the tubes were embedded in casting blocks in groups of five to ten organoids per block, at different timepoints during development for further investigation via immunocytofluorescence. A mixture of 30% sucrose and OCT (50:50) was used for embedding; blocks were stored at -80C until cryosectioning.

### Immunocytochemistry

Using a cryostat (Microm HM 525, Thermo Scientific), the cryopreserved organoids were sectioned at 20 μm thickness and mounted on polarized glass slides. These sections were subjected to heat/citrate-based antigen retrieval for 20 min and permeabilized using Triton X-100 in TBS buffer. The tissue sections were blocked with 10% donkey serum in PBS, 0.1% Triton X-100. Primary antibody [(SOX1, R&D Systems, #AF3369, 1:100), (TBR2, abcam, #ab23345, 1:100), (MAP2, SIGMA, #M9942, 1:200), (ki67, BD Pharmingen, #550609, 1:200), (C-caspase 3, Cell-signaling, #9661S, 1:100) (PCNT, SIGMA, #HPA016820, 1:200), (P-Histone H3, abcam, # ab10543, 1:250), (CTIP2, abcam, #ab18465, 1:500), (SATB2, abcam, #ab51502, 1:150), (HOPX, SIGMA, #HPA030180, 1:500)]incubations were performed at 4°C overnight and secondary incubations were performed at room temperature for 1-3 hr. DAPI staining was performed at 100ng/ml in PBS for 5-7 mins at room temperature. After both primary and secondary antibody incubations, three 20 min washes in PBS-T were performed. AlexaFluor conjugated -anti-mouse 555 (Invitrogen, #A31570), -anti-rabbit 488 (Invitrogen, #A21206), - anti-goat 647 (abcam, #ab150135), -anti-chicken 488 (abcam, #ab150173), -anti-mouse 594 (abcam, #ab150120), -anti-rat 647 (abcam, #ab150167), -anti-rat 568 (Invitrogen, #A11004) were used 1:1000 dilution as secondary antibody. Z-stack images were taken using the confocal microscope (Leica DMi8). Cell counting was performed using “Velocity” software provided by the Imaging Core facility at the Lerner Research Institute, Cleveland Clinic. Violin plots and statistical analysis were generated through GraphPad Prism 9. The images were color enhanced for representation purpose only. The quantification was done using the raw data.

### Neuronal migration assay

We followed the modified protocol from Schaffer et al (2018) (55) for the *in vitro* migration assay, intact organoids cultured for 40 days were resuspended in concentrated Matrigel using wide orifice 200 µL tips and placed on a glass-bottom surface of 4-well culture plates (MatTek; single organoid per well in separate Matrigel drops). The Matrigel was allowed to solidify for 30 min at 37°C, and culture medium was then carefully overlaid on top of Matrigel drops (day 0). Processes were observed emerging from organoids 24 hr later (day 1).

Immunostaining was performed 72 hours post-embedding. For immunostaining, we followed the “whole mount fluorescent immunohistochemistry” provided by Abcam with some modifications. Briefly, the Matrigel-embedded organoids were fixed with 4% paraformaldehyde for 2 hours at room temperature in the fume hood. The tissues were washed and permeabilized with Triton X-100 in PBS. Blocking was performed using 10% FBS (filtered) in PBS with Triton X-100. After washing the blocking solution, the tissue sections on the slides were incubated with the primary antibodies (TUB-ß3, BioLegend, #801202, 1:1000 and DCX, abcam, #ab153668, 1:1000; manufacturer’s suggested dilution concentrations) for 48 hours at 4°C. Post-primary antibody incubation, the slides were washed with Triton X-100 in PBS. The washed sections were then incubated with secondary antibodies (AlexaFluor conjugated) using 1:1000 dilutions and were incubated at 4°C for another 48 hours. Eventually, the tissue sections were counterstained with DAPI for 1 hour at room temperature. Distance (in μm, from the edge of the organoid) of the migrated neurons along the neuronal fibers was assessed using ImageJ Pro software.

### 10X Single-Cell RNA-Seq Analyses

Raw sequencing data from MOPD I and MOPD I-Rescued were processed using the Cell Ranger v3.1.0 (92) pipeline with default parameters. Reads were aligned to the hg19 genome (GRCh37) and downloaded from ENCODE Reference Sequences to generate feature-barcode matrices. Seurat v4.0.3 was used for quality control and secondary analyses (93). For quality control, cells were discarded if they expressed < 300 genes, > 7500 genes, or had a proportion of mapped reads of > 0.2 to the mitochondrial genome. In addition, all genes that were detected in < 5 cells were discarded for downstream analyses. Following log normalization of the filtered matrix, unwanted sources of variations, which include the proportion of mitochondrial transcripts, were regressed out. To improve clustering of samples for downstream analyses, the samples were integrated using canonical correlation analysis (CCA) by selecting the top 3000 most variant genes from each sample. Anchors were identified using mutual nearest neighbors across datasets to integrate the samples. Dimension reduction was performed using PCA on the first 40 principal components followed by organization into clusters using the Louvain algorithm with the Seurat FindClusters function. Downstream cluster visualization was performed via UMAP. Cell type annotation of cerebral organoid cell types was performed using specific markers for each cell type. Differential expression was performed among clusters using the FindMarkers function in Seurat. Pathway enrichment analysis was performed on the DEG sets with the enrichGO and compareCluster functions using the R package clusterProfiler v4.0.5 (94).

Pseudotime trajectory analysis was performed using monocle3 v1.0.0 (95, 96, 97) using default parameters. To specifically analyze the neuronal differentiation pathway in cerebral organoids, we isolated radial glial cells, cycling progenitors, intermediate progenitors, and neurons. This was followed by downsampling each organoid to contain the same number of cells for downstream analysis. Pseudotime trajectory analysis was performed using monocle3 default parameters with the starting node of neuronal differentiation placed on immature radial glial cells. A two-sided Kolmogorov-Smirnov test was performed to test whether MOPD I organoid cell distributions were skewed along the developmental trajectory.

To gain perspective into co-regulated gene expression patterns along the pseudotime trajectory for neuronal differentiation, WGCNA (98, 99) was performed on the downsampled organoids. WGCNA parameters including power threshold were used as defined in Paulsen et al., 2020 (100). A final power of 2 was used for WGCNA analysis, which captures the most variation within the fewest gene expression modules for the MOPD I and MOPD I-Rescued organoids. This resulted in five gene module lists (Supp. Table 2). Each list of genes was incorporated into the organoid datasets using the AddModuleScore function in Seurat. Significance in module expression was determined using a Wilcoxon Rank Sum test.

## Author Contributions

J.S. conceived the project with guidance from F.P., A.W.B., and R.A.P.

All experiments were discussed and designed with the help of R.A.P. and were performed by J.S. Single-cell suspension and capture was performed by J.S., scRNA sequencing data analysis was performed by N.J.D.

J.S. and N.J.D. prepared the figures with significant assistance from R.L.G. The manuscript was prepared with input from all authors.

## Acknowledgments

The authors are grateful to Luke A. Bury and Ya Chen from the A.W.B. lab and Ashleigh Shaffer for their coordination with tissue culture space for generation of organoids. We thank Kimberly Aldinger from Seattle Children’s Hospital for her initial guidance with single-cell RNA sequencing analysis, as well as Judith A. Drazba and Ajay Zalavadia from the Lerner Research Institute Imaging Core facility for their help. We also thank Kewal Asosingh (Director, Flow Core) and Dean Horton (Genoimc Cores) for their initial guidance with 10X Genomics single-cell RNA sequencing. This research project was funded by NIH Grant (R01GM133989) to R.A.P and the 10X Genomics single-cell RNA sequencing experiment was funded by an internal Grant to R.A.P (Q310XPadgett2019).

## Figure Legends

**Supplemental Table 1**

Bulk RNA sequencing from MOPD I iPSCs show intron retention in the genes involved in neurodevelopment. Terminal dinucleotides of these introns and other related details of these genes were extracted from IAOD (Database, Moyer D.C. et al., 2020).

**Supplementary Figure 1.**
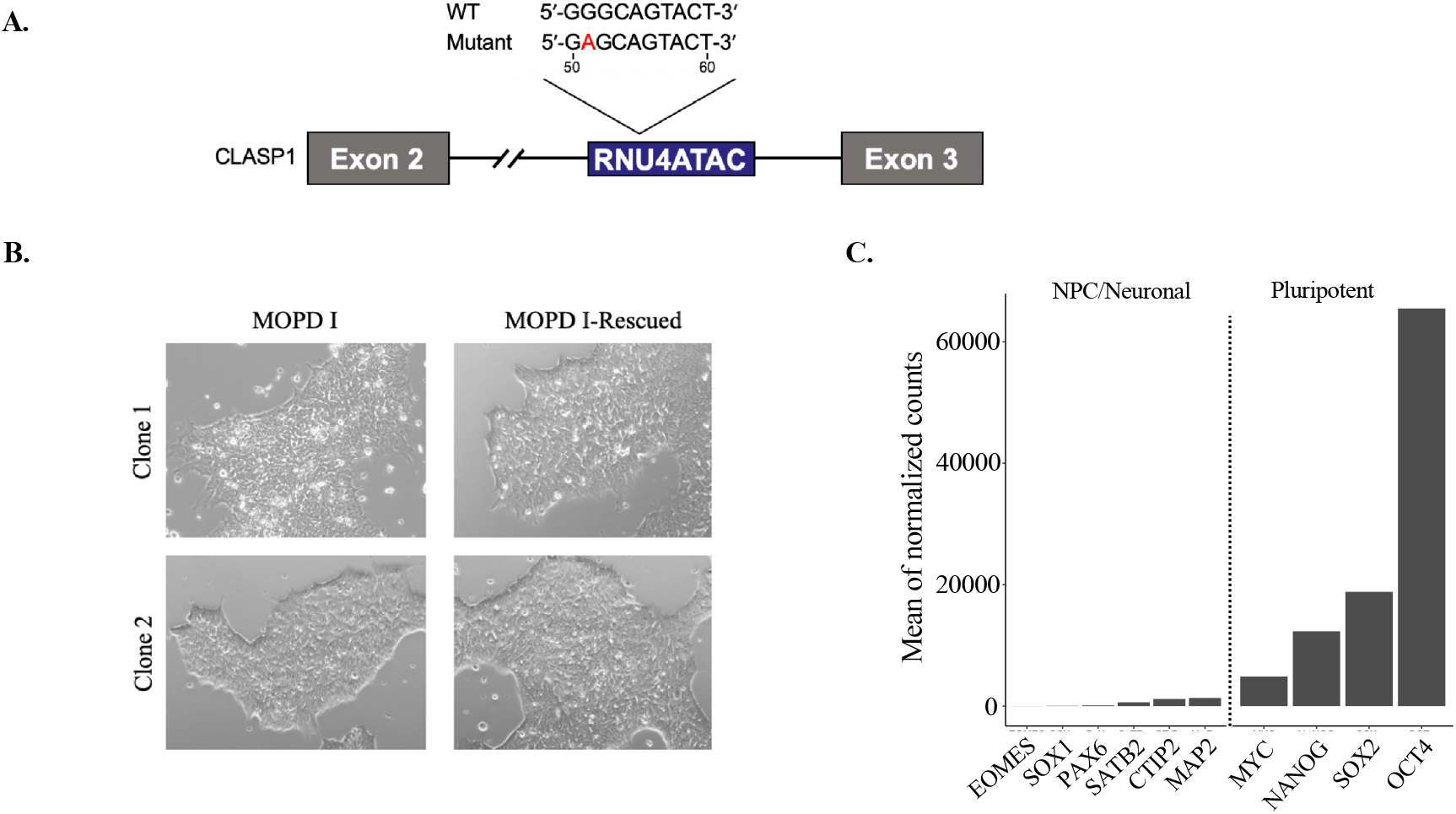
A. Schematics of the CLASP 1 gene. The gray boxes represent exon 2 and exon 3. RNU4ATAC is expressed from the intron 2 (highlighted in blue box) of the CLASP 1 gene. The WT and 51G>A mutant sequences are shown above. B. Representative pictures of iPSCs colonies from MOPD I iPSCs homozygous for the 51G>A mutation and the iPSCs after rescuing with WTU4atac expression. C. Bulk RNA sequencing of iPSC cells showing the expression of pluripotent stem cell markers and neuronal markers. Representative results from the iPSCs clone 1.

**Supplementary Figure 2.**
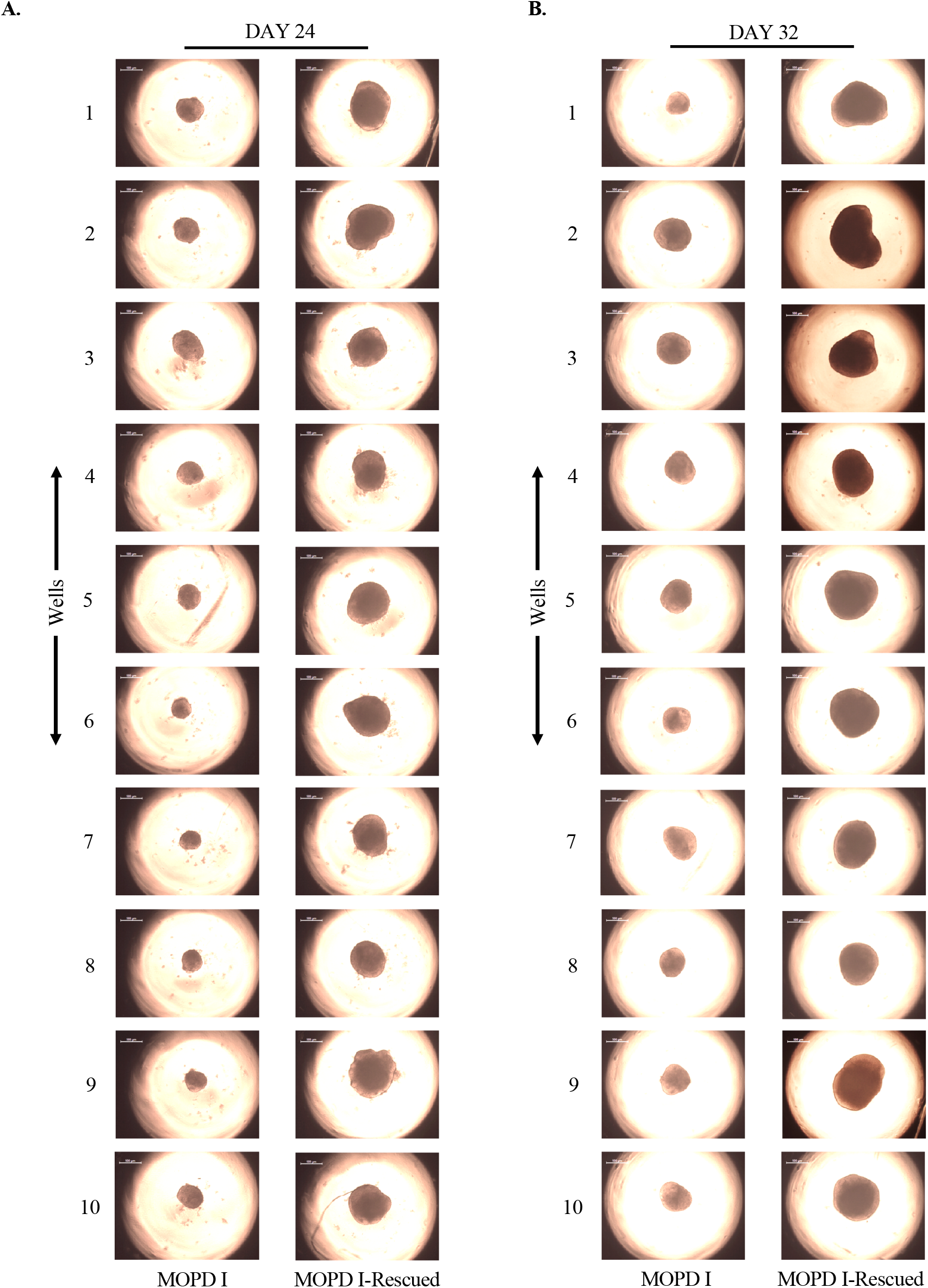
Representative pictures of MOPD I and MOPD I-Rescued forebrain organoids at 24 (left) and 32 (right) days of differentiation.

**Supplementary Figure 3.**
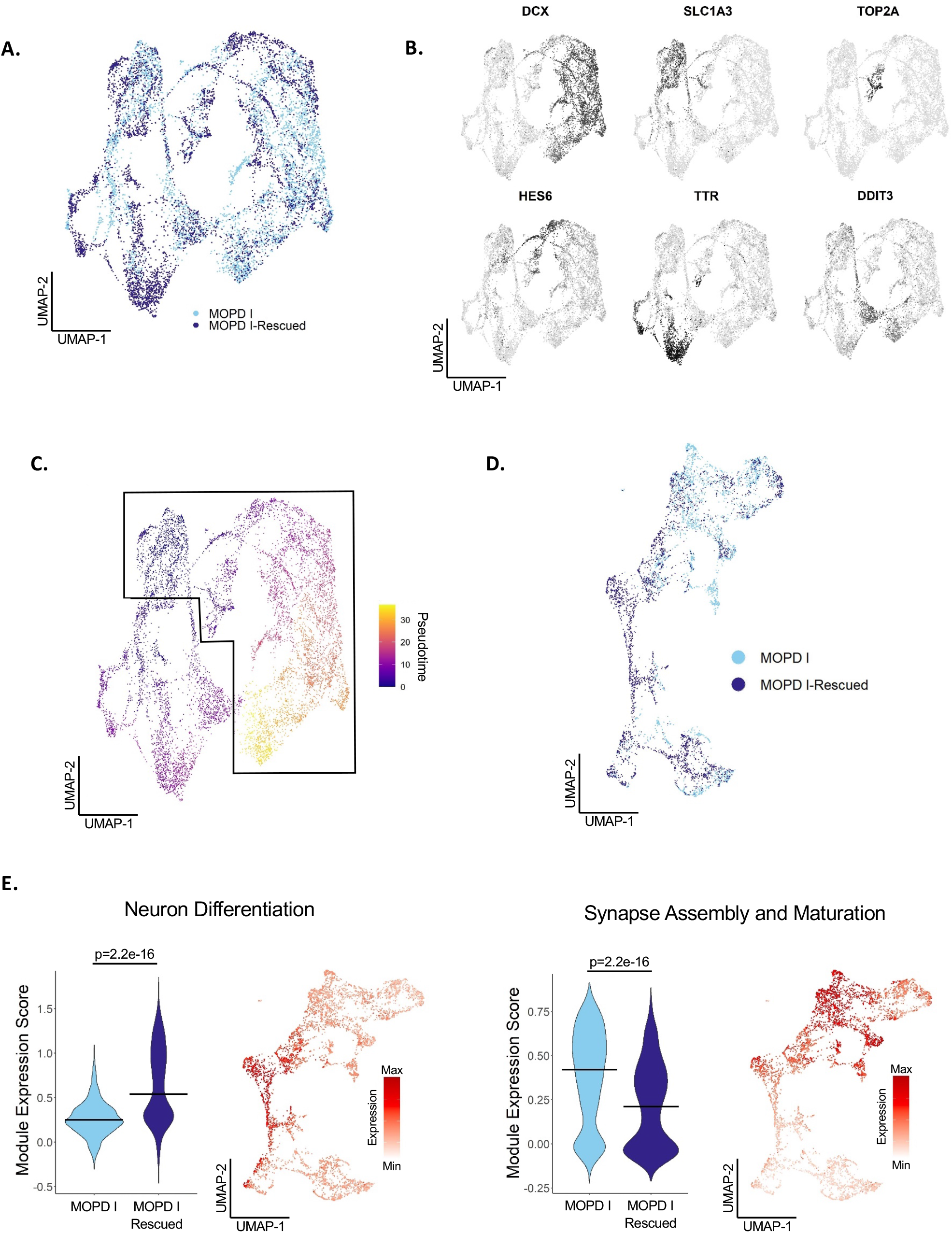
Single-cell RNA-seq of organoids shows reduced expression of genes involved in neuronal differentiation and increased expression of the genes involved in synapse and maturation. A) UMAP of MOPD I and MOPD I-Rescued organoids containing 9454 cells (MOPD I=4448; MOPD I-Rescued=5006). B) UMAP illustrating cell type specific expression markers (DCX/neurons, SLC1A3/radial glial cells, TOP2A/cycling progenitors, HES6/intermediate progenitors, TTR/choroid plexus, DDIT3/UPRC). C) Developmental pseudotime trajectories of MOPD I and MOPD I-Rescued organoids using Monocle3. Starting point of trajectory was placed on early radial glial cells. Cells in the neuronal differentiation pathway are in the black outline and used for subclustering in figure D. D) Subclustering of boxed cells in figure C using Monocle3 and downsampled to include the same number of cells, 2961 cells per sample. E) Module scores of highly correlated gene expression networks in organoids (figure D) identified using WGCNA. Names of modules were defined as relevant pathways using gene ontology analysis of genes enriched in each module. Violin plots depicting the distribution of module scores between MOPD I and MOPD I-Rescued (left). The horizontal bar depicts the median module score for each organoid. UMAP of module scores per cell (right).

**Supplementary Figure 4.**
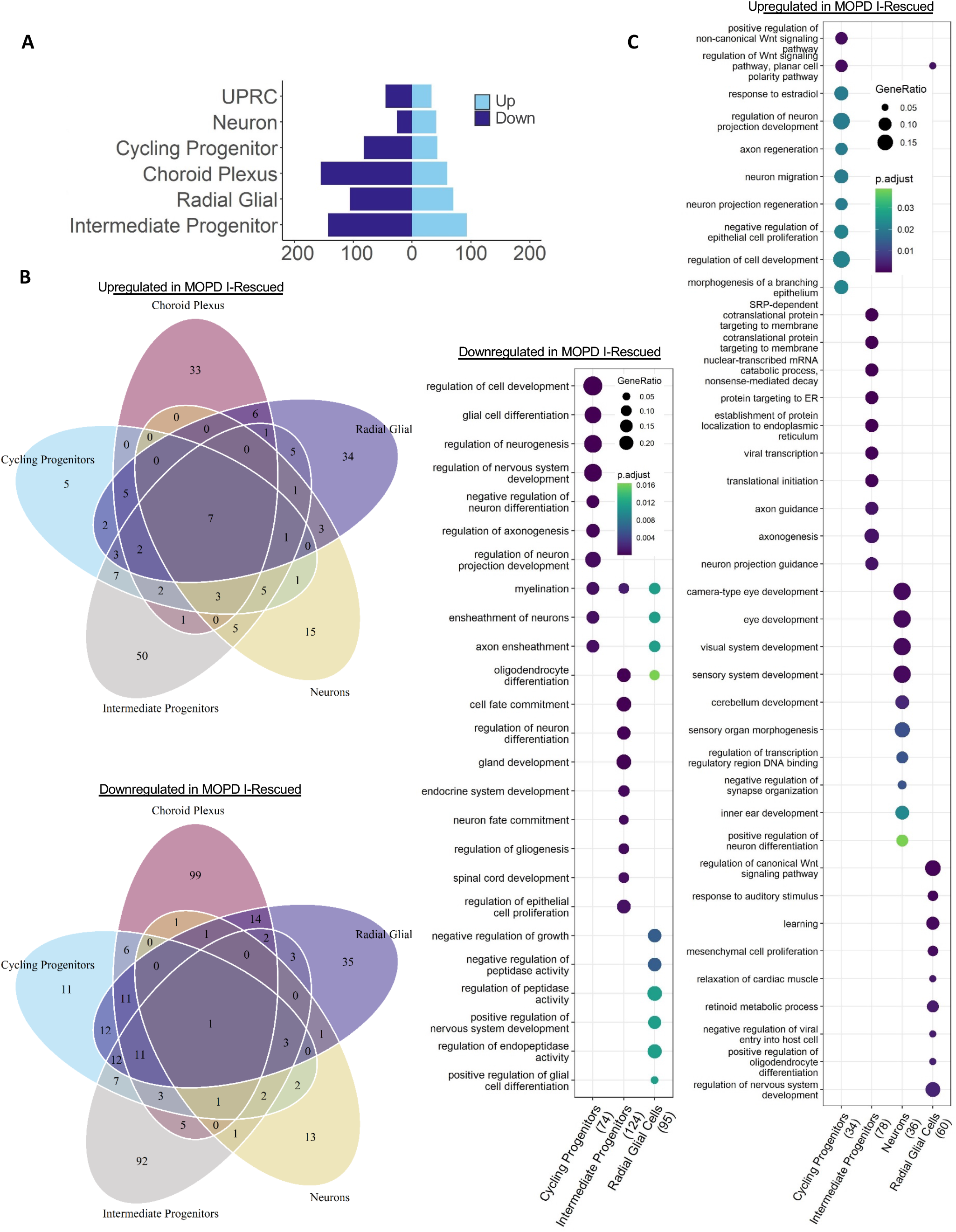
A) Number of significant differentially expressed genes (p value ≤ 0.05, log2FC ≥ 0.75 or ≤ - 0.75) per cell type in MOPD I-Rescued versus MOPD-I. B) Number of common and distinct differentially upregulated (upper panel) and downregulated (lower panel) genes among cell types. Significant genes are from Figure A. C) Gene ontology terms enriched from upregulated (right) or downregulated (left) genes per cell type in MOPD I-Rescued versus MOPD-I. Significant genes were taken from Figure A to run gene ontology enrichment.

**Supplementary Figure 5.**
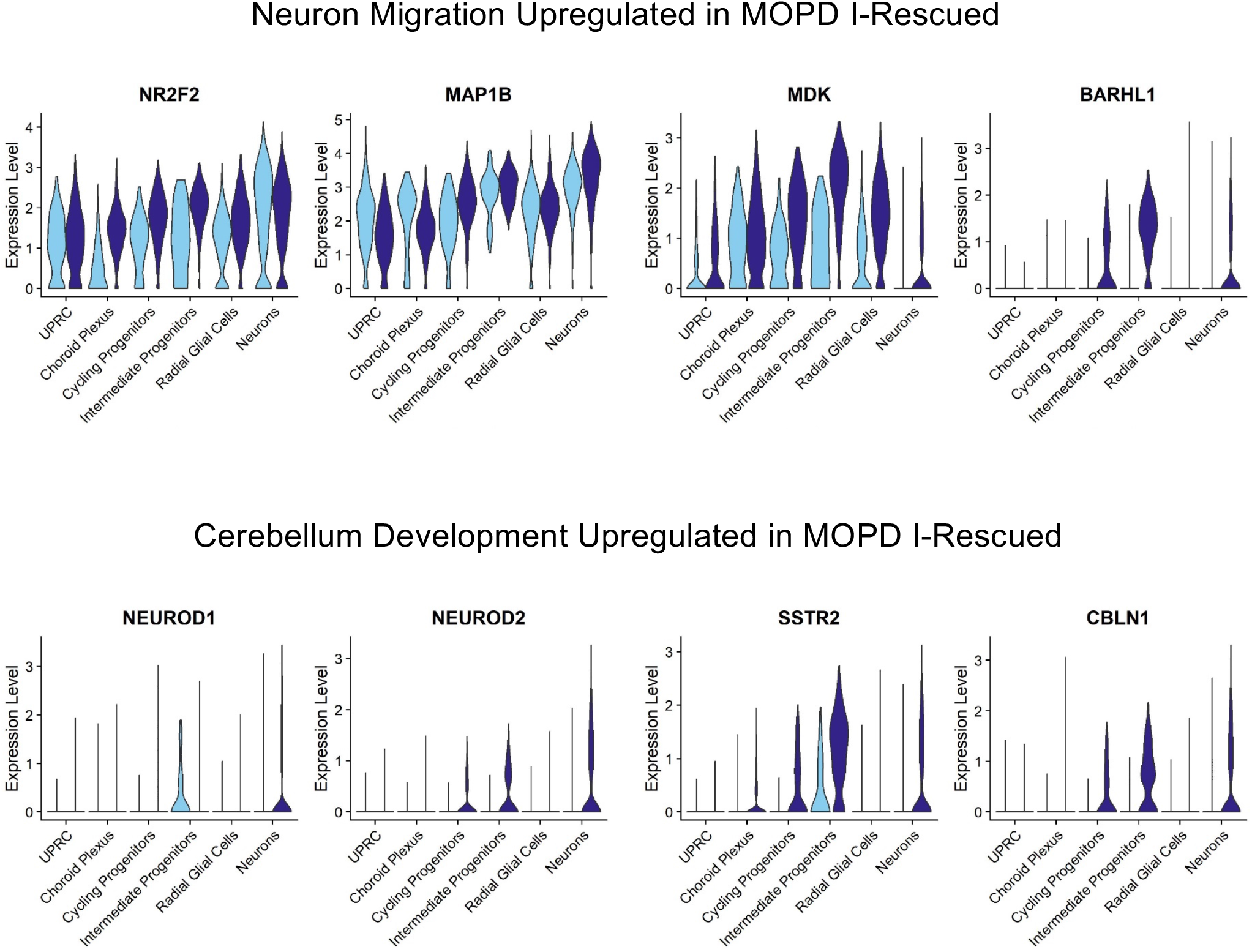
Expression of genes upregulated in enriched GO terms from MOPD I-Rescued versus MOPD I gene ontology analysis.

**Supplementary Table 1.** Increased U12-dependent introns retained in MOPD I iPSCs

Increased U12-dependent introns retained in MOPD I iPSCs
| Gene with retained U12-type intron | Terminal intron dinucleotides | Associated with... |
| --- | --- | --- |
| *TCTN3* | *GT-GG* | Skeletal Dysplasia, agenesis of Corpus Callosum.<br>*Valente, E. M. et al. Nat. Rev. Neurol. 10,27-36 (2014)* |
| *RAF1* | *GT-AG* | Increased apoptosis and abnormal differentiation in Hippocampus resulting in cognitive impairment.<br>*Kim, Y. E. and Baek, S. T. Mol. Cells. 42, 441-447 (2019)* |
| *ZCCHC8* | *AT-AC* | Defects in the development of the central nervous system.<br>*Gable, D. L. et al. Genes & Development. 33, 1381-1396 (2019)* |
| *MAPK12* | *GT-AG* | Regulates function of different glutamate receptors as well as actin dynamics at excitatory synapses.<br>*Mitz, A. R. et al. Euro. J. Hum. Gen. 26, 293-302 (2018)* |

## References

1. Turunen JJ, Niemela EH, Verma B, Frilander MJ. The significant other: splicing by the minor spliceosome. Wiley Interdiscip Rev RNA 4, 61–76 (2013).

2. Moyer DC, Larue GE, Hershberger CE, Roy SW, Padgett RA. Comprehensive database and evolutionary dynamics of U12-type introns. Nucleic Acids Res 48, 7066–7078 (2020).

3. Niemelä EH, Frilander MJ. Regulation of gene expression through inefficient splicing of U12-type introns. RNA Biol 11, 1325–1329 (2014).

4. Verma B, Akinyi MV, Norppa AJ, Frilander MJ. Minor spliceosome and disease. Semin Cell Dev Biol 79, 103–112 (2018).

5. El Marabti E, Malek J, Younis I. Minor Intron Splicing from Basic Science to Disease. Int J Mol Sci 22, (2021).

6. Padgett RA. New connections between splicing and human disease. Trends Genet 28, 147–154 (2012).

7. He H, et al. Mutations in U4atac snRNA, a component of the minor spliceosome, in the developmental disorder MOPD I. Science 332, 238–240 (2011).

8. Edery P, et al. Association of TALS developmental disorder with defect in minor splicing component U4atac snRNA. Science 332, 240–243 (2011).

9. Merico D, et al. Compound heterozygous mutations in the noncoding RNU4ATAC cause Roifman Syndrome by disrupting minor intron splicing. Nat Commun 6, 8718 (2015).

10. Farach LS, et al. The expanding phenotype of RNU4ATAC pathogenic variants to Lowry Wood syndrome. Am J Med Genet A 176, 465–469 (2018).

11. Wang Y, et al. Identification of compound heterozygous variants in the noncoding RNU4ATAC gene in a Chinese family with two successive foetuses with severe microcephaly. Hum Genomics 12, 3 (2018).

12. Nagy R, et al. Microcephalic osteodysplastic primordial dwarfism type I with biallelic mutations in the RNU4ATAC gene. Clin Genet 82, 140–146 (2012).

13. Krøigård AB, Frost M, Larsen MJ, Ousager LB, Frederiksen AL. Bone structure in two adult subjects with impaired minor spliceosome function resulting from RNU4ATAC mutations causing microcephalic osteodysplastic primordial dwarfism type 1 (MOPD1). Bone 92, 145–149 (2016).

14. Otake LR, Scamborova P, Hashimoto C, Steitz JA. The divergent U12-type spliceosome is required for pre-mRNA splicing and is essential for development in Drosophila. Mol Cell 9, 439–446 (2002).

15. König H, Matter N, Bader R, Thiele W, Müller F. Splicing segregation: the minor spliceosome acts outside the nucleus and controls cell proliferation. Cell 131, 718–729 (2007).

16. Kim WY, et al. The Arabidopsis U12-type spliceosomal protein U11/U12-31K is involved in U12 intron splicing via RNA chaperone activity and affects plant development. Plant Cell 22, 3951–3962 (2010).

17. Jung HJ, Kang H. The Arabidopsis U11/U12-65K is an indispensible component of minor spliceosome and plays a crucial role in U12 intron splicing and plant development. Plant J 78, 799–810 (2014).

18. Markmiller S, et al. Minor class splicing shapes the zebrafish transcriptome during development. Proc Natl Acad Sci U S A 111, 3062–3067 (2014).

19. Baumgartner M, et al. Minor splicing snRNAs are enriched in the developing mouse CNS and are crucial for survival of differentiating retinal neurons. Dev Neurobiol 75, 895–907 (2015).

20. Baumgartner M, et al. Minor spliceosome inactivation causes microcephaly, owing to cell cycle defects and death of self-amplifying radial glial cells. Development 145, (2018).

21. Lui JH, Hansen DV, Kriegstein AR. Development and evolution of the human neocortex. Cell 146, 18–36 (2011).

22. Sidman RL, Rakic P. Neuronal migration, with special reference to developing human brain: a review. Brain Res 62, 1–35 (1973).

23. Stiles J, Jernigan TL. The basics of brain development. Neuropsychol Rev 20, 327–348 (2010).

24. Guerrini R, Dobyns WB. Malformations of cortical development: clinical features and genetic causes. Lancet Neurol 13, 710–726 (2014).

25. Hu WF, Chahrour MH, Walsh CA. The diverse genetic landscape of neurodevelopmental disorders. Annu Rev Genomics Hum Genet 15, 195–213 (2014).

26. Homem CC, Repic M, Knoblich JA. Proliferation control in neural stem and progenitor cells. Nat Rev Neurosci 16, 647–659 (2015).

27. Calegari F, Huttner WB. An inhibition of cyclin-dependent kinases that lengthens, but does not arrest, neuroepithelial cell cycle induces premature neurogenesis. J Cell Sci 116, 4947–4955 (2003).

28. Lange C, Huttner WB, Calegari F. Cdk4/cyclinD1 overexpression in neural stem cells shortens G1, delays neurogenesis, and promotes the generation and expansion of basal progenitors. Cell Stem Cell 5, 320–331 (2009).

29. Pilaz LJ, et al. Prolonged Mitosis of Neural Progenitors Alters Cell Fate in the Developing Brain. Neuron 89, 83–99 (2016).

30. Pilaz LJ, et al. Forced G1-phase reduction alters mode of division, neuron number, and laminar phenotype in the cerebral cortex. Proc Natl Acad Sci U S A 106, 21924–21929 (2009).

31. Kowalczyk T, et al. Intermediate neuronal progenitors (basal progenitors) produce pyramidal-projection neurons for all layers of cerebral cortex. Cereb Cortex 19, 2439–2450 (2009).

32. Bershteyn M, et al. Human iPSC-Derived Cerebral Organoids Model Cellular Features of Lissencephaly and Reveal Prolonged Mitosis of Outer Radial Glia. Cell Stem Cell 20, 435-449.e434 (2017).

33. Jafarifar F, Dietrich RC, Hiznay JM, Padgett RA. Biochemical defects in minor spliceosome function in the developmental disorder MOPD I. Rna 20, 1078–1089 (2014).

34. Hodge, RD, Bakken, TE, Miller, JA et al. Conserved cell types with divergent features in human versus mouse cortex. Nature 573, 61–68 (2019).

35. Arlotta P, Paşca SP. Cell diversity in the human cerebral cortex: from the embryo to brain organoids. Curr Opin Neurobiol 56, 194–198 (2019).

36. Quadrato G, et al. Cell diversity and network dynamics in photosensitive human brain organoids. Nature 545, 48–53 (2017).

37. Seto Y, Eiraku M. Human brain development and its in vitro recapitulation. Neurosci Res 138, 33–42 (2019).

38. Velasco S, et al. Individual brain organoids reproducibly form cell diversity of the human cerebral cortex. Nature 570, 523–527 (2019).

39. Kadoshima T, et al. Self-organization of axial polarity, inside-out layer pattern, and species-specific progenitor dynamics in human ES cell-derived neocortex. Proc Natl Acad Sci U S A 110, 20284–20289 (2013).

40. Fish JL, Kosodo Y, Enard W, Pääbo S, Huttner WB. Aspm specifically maintains symmetric proliferative divisions of neuroepithelial cells. Proc Natl Acad Sci U S A 103, 10438–10443 (2006).

41. Yingling J, et al. Neuroepithelial stem cell proliferation requires LIS1 for precise spindle orientation and symmetric division. Cell 132, 474–486 (2008).

42. Pawlisz AS, Mutch C, Wynshaw-Boris A, Chenn A, Walsh CA, Feng Y. Lis1-Nde1-dependent neuronal fate control determines cerebral cortical size and lamination. Hum Mol Genet 17, 2441–2455 (2008).

43. Pramparo T, Youn YH, Yingling J, Hirotsune S, Wynshaw-Boris A. Novel embryonic neuronal migration and proliferation defects in Dcx mutant mice are exacerbated by Lis1 reduction. J Neurosci 30, 3002–3012 (2010).

44. Xie Y, Jüschke C, Esk C, Hirotsune S, Knoblich JA. The phosphatase PP4c controls spindle orientation to maintain proliferative symmetric divisions in the developing neocortex. Neuron 79, 254–265 (2013).

45. Pirozzi F, Nelson B, Mirzaa G. From microcephaly to megalencephaly: determinants of brain size. Dialogues Clin Neurosci 20, 267–282 (2018).

46. Chowdhury R, et al. STAU2 binds a complex RNA cargo that changes temporally with production of diverse intermediate progenitor cells during mouse corticogenesis. Development 148, (2021).

47. Molyneaux BJ, Arlotta P, Hirata T, Hibi M, Macklis JD. Fezl is required for the birth and specification of corticospinal motor neurons. Neuron 47, 817–831 (2005).

48. Leone DP, Srinivasan K, Chen B, Alcamo E, McConnell SK. The determination of projection neuron identity in the developing cerebral cortex. Curr Opin Neurobiol 18, 28–35 (2008).

49. Hirotsune S, et al. Graded reduction of Pafah1b1 (Lis1) activity results in neuronal migration defects and early embryonic lethality. Nat Genet 19, 333–339 (1998).

50. Gambello MJ, Darling DL, Yingling J, Tanaka T, Gleeson JG, Wynshaw-Boris A. Multiple dose-dependent effects of Lis1 on cerebral cortical development. J Neurosci 23, 1719–1729 (2003).

51. Youn YH, Pramparo T, Hirotsune S, Wynshaw-Boris A. Distinct dose-dependent cortical neuronal migration and neurite extension defects in Lis1 and Ndel1 mutant mice. J Neurosci 29, 15520–15530 (2009).

52. Toyo-oka K, et al. 14-3-3epsilon is important for neuronal migration by binding to NUDEL: a molecular explanation for Miller-Dieker syndrome. Nat Genet 34, 274–285 (2003).

53. Sheen VL, et al. Neocortical neuronal arrangement in Miller Dieker syndrome. Acta Neuropathol 111, 489–496 (2006).

54. Saito T, et al. Neocortical layer formation of human developing brains and lissencephalies: consideration of layer-specific marker expression. Cereb Cortex 21, 588–596 (2011).

55. Schaffer AE, et al. Biallelic loss of human CTNNA2, encoding αN-catenin, leads to ARP2/3 complex overactivity and disordered cortical neuronal migration. Nat Genet 50, 1093–1101 (2018).

56. Gai M, et al. ASPM and CITK regulate spindle orientation by affecting the dynamics of astral microtubules. EMBO Rep 17, 1396–1409 (2016).

57. Higgins J, et al. Human ASPM participates in spindle organisation, spindle orientation and cytokinesis. BMC Cell Biol 11, 85 (2010).

58. Tungadi EA, Ito A, Kiyomitsu T, Goshima G. Human microcephaly ASPM protein is a spindle pole-focusing factor that functions redundantly with CDK5RAP2. J Cell Sci 130, 3676–3684 (2017).

59. Verloes A, Drunat S, Passemard S. ASPM Primary Microcephaly. In: GeneReviews(®) (eds Adam MP, et al.). University of Washington, Seattle Copyright © 1993-2022, University of Washington, Seattle. GeneReviews is a registered trademark of the University of Washington, Seattle. All rights reserved. (1993).

60. Belvindrah R, et al. Mutation of the α-tubulin Tuba1a leads to straighter microtubules and perturbs neuronal migration. J Cell Biol 216, 2443–2461 (2017).

61. Fernández-Santiago R, Ezquerra M. Epigenetic Research of Neurodegenerative Disorders Using Patient iPSC-Based Models. Stem Cells Int 2016, 9464591 (2016).

62. Grainger AI, King MC, Nagel DA, Parri HR, Coleman MD, Hill EJ. In vitro Models for Seizure-Liability Testing Using Induced Pluripotent Stem Cells. Front Neurosci 12, 590 (2018).

63. Hattori N. Cerebral organoids model human brain development and microcephaly. Mov Disord 29, 185 (2014).

64. Ishii MN, Yamamoto K, Shoji M, Asami A, Kawamata Y. Human induced pluripotent stem cell (hiPSC)-derived neurons respond to convulsant drugs when co-cultured with hiPSC-derived astrocytes. Toxicology 389, 130–138 (2017).

65. Kaufmann M, et al. High-Throughput Screening Using iPSC-Derived Neuronal Progenitors to Identify Compounds Counteracting Epigenetic Gene Silencing in Fragile X Syndrome. J Biomol Screen 20, 1101–1111 (2015).

66. Li R, Sun L, Fang A, Li P, Wu Q, Wang X. Recapitulating cortical development with organoid culture in vitro and modeling abnormal spindle-like (ASPM related primary) microcephaly disease. Protein Cell 8, 823–833 (2017).

67. Liang N, Trujillo CA, Negraes PD, Muotri AR, Lameu C, Ulrich H. Stem cell contributions to neurological disease modeling and personalized medicine. Prog Neuropsychopharmacol Biol Psychiatry 80, 54–62 (2018).

68. McMillan HJ, et al. Whole genome sequencing identifies pathogenic RNU4ATAC variants in a child with recurrent encephalitis, microcephaly, and normal stature. Am J Med Genet A 185, 3502–3506 (2021).

69. Nevin ZS, et al. Modeling the Mutational and Phenotypic Landscapes of Pelizaeus-Merzbacher Disease with Human iPSC-Derived Oligodendrocytes. Am J Hum Genet 100, 617–634 (2017).

70. Shi Y, Inoue H, Wu JC, Yamanaka S. Induced pluripotent stem cell technology: a decade of progress. Nat Rev Drug Discov 16, 115–130 (2017).

71. Tang BL. Patient-Derived iPSCs and iNs-Shedding New Light on the Cellular Etiology of Neurodegenerative Diseases. Cells 7, (2018).

72. Tilgner K, et al. A human iPSC model of Ligase IV deficiency reveals an important role for NHEJ-mediated-DSB repair in the survival and genomic stability of induced pluripotent stem cells and emerging haematopoietic progenitors. Cell Death Differ 20, 1089–1100 (2013).

73. Zhang M, Ngo J, Pirozzi F, Sun YP, Wynshaw-Boris A. Highly efficient methods to obtain homogeneous dorsal neural progenitor cells from human and mouse embryonic stem cells and induced pluripotent stem cells. Stem Cell Res Ther 9, 67 (2018).

74. Wong M, Roper SN. Genetic animal models of malformations of cortical development and epilepsy. J Neurosci Methods 260, 73–82 (2016).

75. Lancaster MA, et al. Cerebral organoids model human brain development and microcephaly. Nature 501, 373–379 (2013).

76. Neumüller RA, Knoblich JA. Dividing cellular asymmetry: asymmetric cell division and its implications for stem cells and cancer. Genes Dev 23, 2675–2699 (2009).

77. Siller KH, Doe CQ. Spindle orientation during asymmetric cell division. Nat Cell Biol 11, 365–374 (2009).

78. Morin X, Bellaïche Y. Mitotic spindle orientation in asymmetric and symmetric cell divisions during animal development. Dev Cell 21, 102–119 (2011).

79. Faheem M, et al. Molecular genetics of human primary microcephaly: an overview. BMC Med Genomics 8 Suppl 1, S4 (2015).

80. Farag HG, et al. Abnormal centrosome and spindle morphology in a patient with autosomal recessive primary microcephaly type 2 due to compound heterozygous WDR62 gene mutation. Orphanet J Rare Dis 8, 178 (2013).

81. Issa L, et al. CDK5RAP2 expression during murine and human brain development correlates with pathology in primary autosomal recessive microcephaly. Cereb Cortex 23, 2245–2260 (2013).

82. Moon HM, Youn YH, Pemble H, Yingling J, Wittmann T, Wynshaw-Boris A. LIS1 controls mitosis and mitotic spindle organization via the LIS1-NDEL1-dynein complex. Hum Mol Genet 23, 449–466 (2014).

83. Nicholas AK, et al. WDR62 is associated with the spindle pole and is mutated in human microcephaly. Nat Genet 42, 1010–1014 (2010).

84. Noctor SC, Flint AC, Weissman TA, Dammerman RS, Kriegstein AR. Neurons derived from radial glial cells establish radial units in neocortex. Nature 409, 714–720 (2001).

85. Noctor SC, Martínez-Cerdeño V, Ivic L, Kriegstein AR. Cortical neurons arise in symmetric and asymmetric division zones and migrate through specific phases. Nat Neurosci 7, 136–144 (2004).

86. Pontious A, Kowalczyk T, Englund C, Hevner RF. Role of intermediate progenitor cells in cerebral cortex development. Dev Neurosci 30, 24–32 (2008).

87. Létard P, et al. Autosomal recessive primary microcephaly due to ASPM mutations: An update. Hum Mutat 39, 319–332 (2018).

88. Hong SE, et al. Autosomal recessive lissencephaly with cerebellar hypoplasia is associated with human RELN mutations. Nat Genet 26, 93–96 (2000).

89. Camp JG, et al. Human cerebral organoids recapitulate gene expression programs of fetal neocortex development. Proc Natl Acad Sci U S A 112, 15672–15677 (2015).

90. Poirier K, et al. Large spectrum of lissencephaly and pachygyria phenotypes resulting from de novo missense mutations in tubulin alpha 1A (TUBA1A). Hum Mutat 28, 1055–1064 (2007).

91. Zhou W, Freed CR. Adenoviral gene delivery can reprogram human fibroblasts to induced pluripotent stem cells. Stem Cells 27, 2667–2674 (2009).

92. Zheng GX, et al. Massively parallel digital transcriptional profiling of single cells. Nat Commun 8, 14049 (2017).

93. Hao Y, et al. Integrated analysis of multimodal single-cell data. Cell 184, 3573-3587.e3529 (2021).

94. Yu G, Wang LG, Han Y, He QY. clusterProfiler: an R package for comparing biological themes among gene clusters. Omics 16, 284–287 (2012).

95. Trapnell C, et al. The dynamics and regulators of cell fate decisions are revealed by pseudotemporal ordering of single cells. Nat Biotechnol 32, 381–386 (2014).

96. Qiu X, et al. Reversed graph embedding resolves complex single-cell trajectories. Nat Methods 14, 979–982 (2017).

97. Cao J, et al. The single-cell transcriptional landscape of mammalian organogenesis. Nature 566, 496–502 (2019).

98. Zhang B, Horvath S. A general framework for weighted gene co-expression network analysis. Stat Appl Genet Mol Biol 4, Article17 (2005).

99. Langfelder P, Horvath S. WGCNA: an R package for weighted correlation network analysis. BMC Bioinformatics 9, 559 (2008).

100. Paulsen B, et al. Human brain organoids reveal accelerated development of cortical neuron classes as a shared feature of autism risk genes. bioRxiv, 2020.2011.2010.376509 (2020).

